# Neural Synchrony Links Sensorimotor Cortices in a Network for Facial Motor Control

**DOI:** 10.1101/2025.03.04.641458

**Authors:** Yuriria Vázquez, Geena R. Ianni, Elie Rassi, Adam G. Rouse, Marc H. Schieber, Faraz Yazdani, Yifat Prut, Winrich A. Freiwald

## Abstract

Primate societies rely on the production and interpretation of social signals, in particular those displayed by the face. Facial movements are controlled, according to the dominant neuropsychological schema, by two separate circuits, one originating in medial frontal cortex controlling emotional expressions, and a second one originating in lateral motor and premotor areas controlling voluntary facial movements. Despite this functional dichotomy, cortical anatomy suggests that medial and lateral areas are directly connected and may thus operate as a single network. Here we test these contrasting hypotheses through structural and functional magnetic resonance imaging (fMRI) guided electrical stimulation and simultaneous multi-channel recordings from key facial motor areas in the macaque monkey brain. These areas include medial facial motor area M3 (located in the anterior cingulate cortex); two lateral face-related motor areas: M1 (primary motor) and PMv (ventrolateral premotor); and S1 (primary somatosensory cortex). Cortical responses evoked by intracortical stimulation revealed that medial and lateral areas can exert significant functional impact on each other. Simultaneous recordings of local field potentials in all facial motor areas further confirm that during facial expressions, medial and lateral facial motor areas significantly interact, primarily in the alpha and beta frequency ranges, whereas during voluntary chewing, coupling occurs at lower frequencies. These functional interactions varied across facial movement types. Thus, at the cortical level, the control of facial movements is not mediated through independent (medial/lateral) functional streams, but results from an interacting sensorimotor network.

**Significance Statement:** Primates communicate through facial expressions. How the brain generates facial expressions remains poorly understood. To uncover how facial motor-related cortical brain regions interact to produce facial gestures, we combined fMRI-targeted electrophysiology and intracortical microstimulation while monkeys produced qualitatively different facial movements. Our two-pronged experimental approach revealed that facial motor-related cortical areas form an interconnected network characterized by synchronized neural activity demonstrating dynamic expression-selective activity states that are coordinated across the network nodes. Thus, the multiple facial motor-related cortical areas sending axons directly into the facial nucleus operate as a single network in which the overall complex, behavior-specific inter-areal interactions dictate the relevant motor output.

## Introduction

In primate societies, individuals thrive with their ability to interpret and produce social signals^1^. Primates communicate socially through vocalizations, speech, and facial expressions, all of which require precise control of facial musculature ^2,3^. Facial expressions convey information about internal states or intentions, and thereby function as social signals among conspecifics. Despite the significance of facial expressions in social interactions, little is understood about the neural mechanisms underlying these social motor behaviors when compared to other motor actions such as reaching or grasping ^4,5,6,7^.

The primate brain harbors several cortical regions dedicated to facial motor control within the frontal cortex –specifically, the primary motor (M1) and the ventrolateral premotor (PMv) cortices on the lateral aspect, as well as the supplementary (M2) and the cingulate (rostral M3 and caudal M4) motor cortices on the medial aspect ^8,9,10^. Each of these cortical areas project directly to the facial motor nucleus in the brain stem^11^, suggesting that motor cortical areas exert direct control over facial motoneurons in a seemingly parallel, independent control strategy, where each cortical area directly controls a subset of facial muscles.

Studies in human patients have shed light on the functional circuitry of these areas during facial expressions. Two facial areas (M1 and PMv) reside laterally, in the territory of the middle cerebral artery, and three areas (SMA, rostral and caudal cingulate cortex) reside medially, in the territory of the anterior cerebral artery^8^. Patients with a middle cerebral artery stroke display asymmetry in voluntary or goal-directed facial movements (performed in response to verbal commands), yet their ability to perform spontaneous emotional gestures like smiling remains intact^12^. In contrast, patients with an anterior cerebral artery stroke (affecting medial motor areas) can execute normal voluntary facial movements, but struggle with emotional facial gestures^12^. These findings led to the neuropsychological schema that facial movements are controlled by two distinct pathways: lateral and medial motor pathways (Fig.1A) ^13,14^, each controlling a qualitatively different type of facial movement. This is further supported by the unique input and output connectivity patterns of the medial and lateral motor areas of the face. On the other hand, anatomical studies in monkeys have identified corticocortical connectivity between medial and lateral motor areas of the face^11,15,16,17,18,19^, including the facial representation in the somatosensory cortex^8,18^. Brain-wide imaging studies in non-human primates using network connectivity analysis revealed a distributed face motor network whose activation during a specific communicative facial expression extends beyond the medial cingulate cortex to include lateral motor areas^20^.

**Figure 1.**
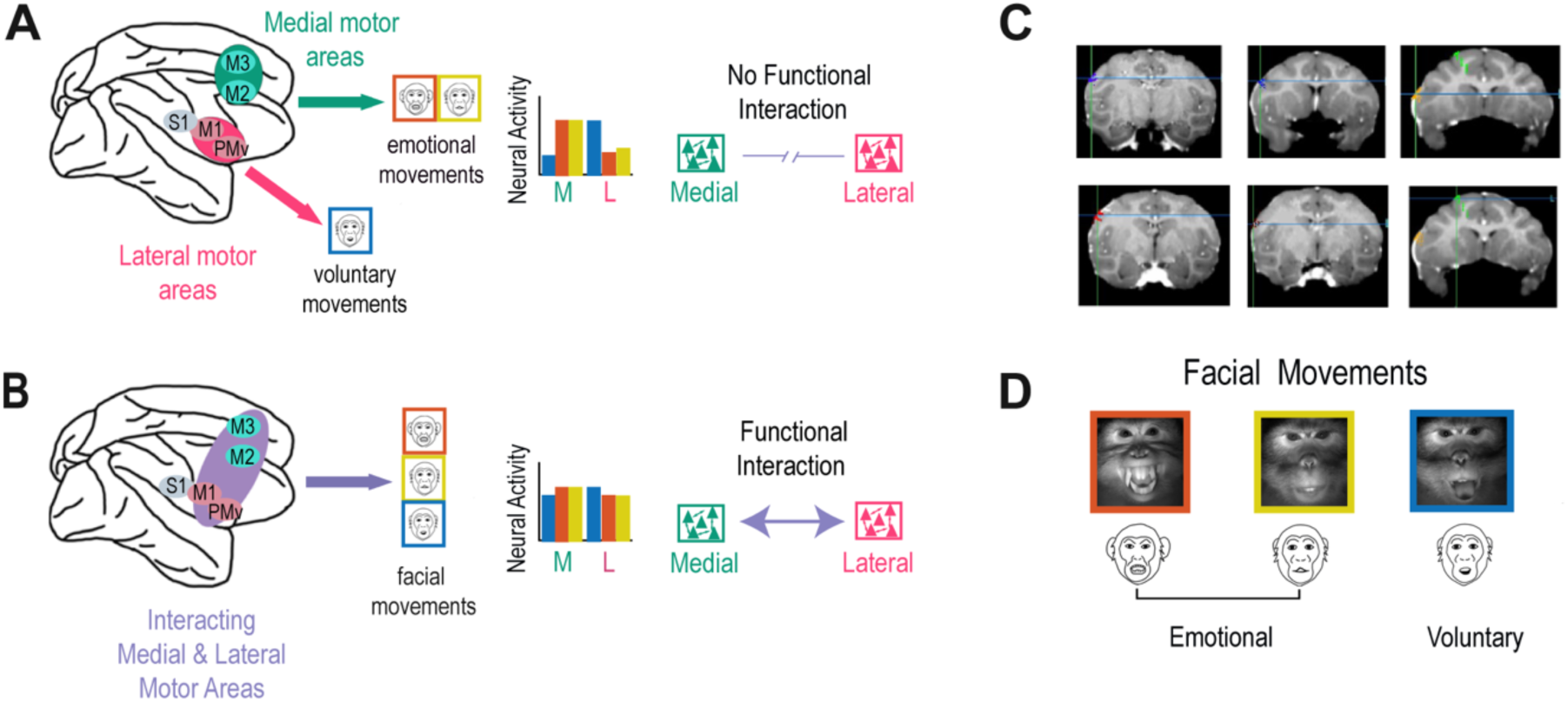
Cortical facial motor representations in the primate brain and facial motor behaviors. Medial (green) and lateral (pink) motor systems are hypothesized to support emotional and voluntary facial movements, respectively. Two possible neural mechanisms for facial movements are proposed: (**A**) Null hypothesis: medial and lateral motor areas operate independently, with medial regions specialized in emotional movements-facial expressions, and lateral areas for voluntary movements such as chewing. (**B**) Alternative hypothesis: medial and lateral motor areas functionally interact to support both emotional and voluntary movements. (**C**) Structural MRI (T1, coronal plane) showing electrode array placement in one monkey. Floating microarrays targeted primary somatosensory (S1, purple), motor (M1, light & dark red) cortex, ventrolateral premotor (PMv, blue & yellow) and cingulate motor cortex (M3, green). Array locations were based on fMRI conjunction maps of facial expressions and mouth movement (see methods). **(D)** Examples of naturalistic facial movements: threat (T, red), a non-affiliative emotional display with sustained open mouth; lip-smack (LS, yellow), a rhythmic affiliative emotional movement of jaw and lips; and chewing (C, blue), an ingestive voluntary movement. Corresponding cartoons below each photo are used as references through the text. Neural activity was recorded simultaneously across all areas during these behaviors, with emotional movements elicited through videos, an avatar and primate interactions (see methods).

Thus, there are two opposing views on the organization of cortical face-motor control: the two parallel pathway hypothesis and the interconnected network hypothesis. The first predicts that there are no functional interactions between medial and lateral facial motor areas. The second hypothesis predicts that such interactions should exist, as medial and lateral cortical areas jointly determine facial movements (Fig. 1B). To test these predictions, we implanted recording arrays in fMRI-identified cortical areas of the facial motor and somatosensory areas of non-human primates. We first determine the capacity of an area to exert an influence on activity in another area by delivering intracortical microstimulation (ICMS) to one brain area while simultaneously recording the evoked local field potentials (LFPs) in the other areas, and estimating the average stimulus triggered response. We then determined the functional interactions taking place during naturalistic behavior, by measuring synchronous patterns in the local field potential (LFPs) among the multiple facial motor areas and the primary somatosensory cortex (S1) to assess functional connectivity.

Our findings reveal that cortical motor areas of the face function as an interconnected network where medial and lateral areas can exert significant influence on each other, as shown through our ICMS approach. During emotional facial expressions, we observed alpha and beta synchronization between medial motor area M3, lateral areas (M1 and PMv) and the face representation in primary somatosensory cortex, suggesting the presence of dynamic, expression-selective activity states coordinated across network nodes. In contrast, the coupling between medial and lateral motor areas during voluntary chewing occurred at lower frequencies, highlighting the frequency-specific nature of cortical coordination patterns for different facial movements. The interareal dynamics of this sensorimotor network displayed different patterns of functional connectivity that correlated with specific facial gestures. This supports our hypothesis that facial motor areas and S1 form an interconnected network with flexible interactions based on synchronized neural activity, generating distinct neural patterns for different types of facial motor behavior.

## Results

We recorded LFPs from multiple channels (n=32 channels per area) located in the sensorimotor areas of the face, including the primary somatosensory cortex (S1), primary motor cortex (M1), ventrolateral premotor cortex (PMv) and cingulate motor cortex (M3), as head-fixed monkeys produced ethologically relevant facial movements in response to various stimuli (see Methods, Fig. 1C-D and Fig. S1).

### Estimating interareal connectivity in the facial motor system with intracortical microstimulation

To identify connectivity between facial motor areas, we employed intracortical microstimulation (ICMS) while the animal was sitting quietly, making no facial movements. For this purpose, we injected single pulse stimuli (250 µA, a train of biphasic pulses, each pulse duration=0.4 ms at a frequency of 3 Hz (333ms inter-pulse interval) delivered for ∼2min) through dedicated low-impedance stimulation electrodes embedded in the recording arrays and estimated the average stimulus triggered response recorded in neighboring arrays (Fig. 2A).

**Figure 2.**
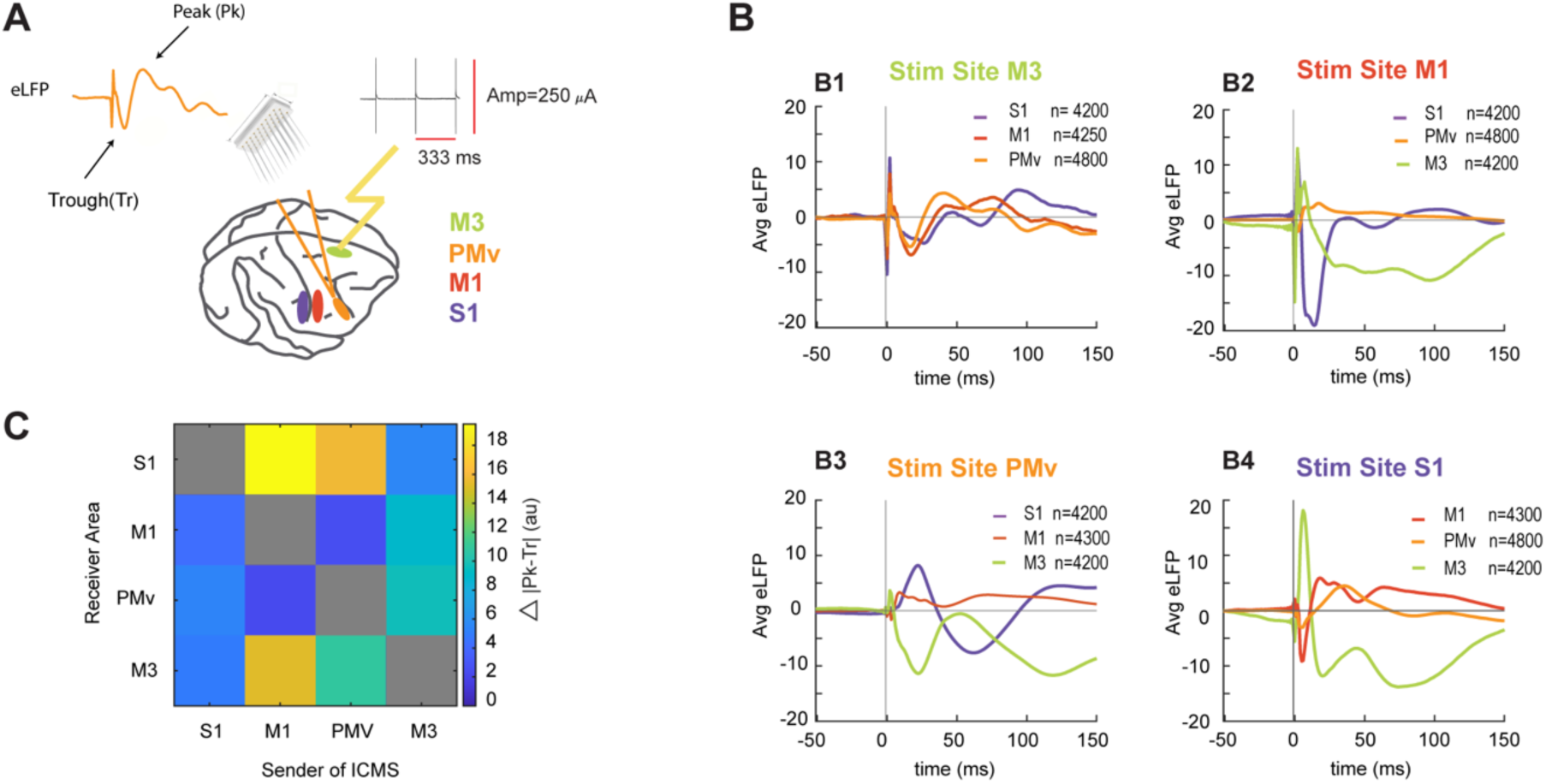
Mapping Facial Sensory and Motor Connectivity through Intracortical Microstimulation. (**A**) Illustration depicting a monkey’s brain with electrode arrays in the recorded areas. The intracortical microstimulation (ICMS) current protocol was applied to the sender area, in this case the M3 cingulate motor area. The protocol uses a train of biphasic pulses (250 µA, each pulse duration=0.4 ms at a frequency of 3 Hz (333 ms inter-pulse interval) delivered for ∼2min). The orange trace represents the PMv’s mean evoked local field potential due to current propagation, highlighting the first peak and trough after the stimulation onset. (**B**) Averaged evoked local field potentials (LFPs) across cortical areas in response to microstimulation of M3 (B1), M1(B2), PMv(B3) and S1 (B4) cortices. The stimulation onset is marked at zero on the x-axis. A stimulation artifact with rapid onset and large amplitude is visible across all averages during the first few milliseconds following stimulation. To exclude this artifact from the analysis, peaks and troughs were searched for starting 6 ms after stimulation onset in all receiving areas. (**C**) Confusion matrix depicting connectivity indexes measured by the absolute difference between the first peak or trough, and the second trough or peak, respectively (Δ|Peak-Trough|). Current sender areas are shown on the *X*-axis and receiver areas on the *Y*-axis. Gray diagonal elements represent untested within-area connectivity, while off-diagonal elements show the measured connectivity strength.

Net signal propagation was assessed by analyzing the eLFP measured in the receiving areas. A key measurement used here was the response magnitude, calculated as the difference between either the first peak and trough or the first trough and peak following the stimulation pulse. To avoid measuring stimulation artifacts, we searched for peaks and troughs starting 6 ms after stimulation onset in all areas, when artifactual deflections had ended.

Figure 2B shows the average area-specific eLFP obtained in response to microstimulation applied in M3 (B1), M1 (B2), PMv (B3) and S1 (B4) obtained across several recording sessions (n ≥ 4200 stimulation pulses). We found evidence for signal propagating from the medial (M3) to the lateral (M1 and PMv) motor areas, and the primary somatosensory cortex (S1). The eLFP responses obtained in S1 (purple line) were generally smaller in amplitude and delayed by ∼10 ms relative to the ones in the lateral motor areas (red and orange lines) suggesting a slower signal propagation to this region (Fig. 2B1). Altogether, we found that ICMS produced significant eLFP responses in all combinations of areas, though with varying magnitudes and waveform patterns. We quantified the evoked connectivity between brain regions using a connectivity matrix (Fig. 2C). This matrix revealed several key patterns, notably that M1 and S1 showed reciprocal connectivity^21^. M3 received inputs from both lateral motor areas (M1 and PMv), with M1 eliciting a larger response than PMv. The response to primary somatosensory (S1) stimulation was comparatively smaller. When stimulation was applied to M3, both lateral motor areas responded (M1 and PMV), aligning with previously described anatomically connectivity patterns^8^. However, an asymmetry was observed: while M3 stimulation produced relatively modest responses in all other areas, stimulation of M1 evoked strong responses in M3. This asymmetry could potentially be explained by orthodromic activation of the source area during local stimulation, supporting M3 receiving inputs from M1, as previously reported^19^.

This set of results validates the use of eLFP for identifying anatomically relevant functional connectivity across different cortical areas and suggests that the observed medial-to-lateral connectivity reveals physiologically relevant pathways between these network components.

### LFP Synchrony Links Sensorimotor Cortices in a Network for Facial Motor Control

We assessed functional connectivity by calculating the synchronization of LFP signals during different facial gestures in monkeys. This approach allowed us to examine how brain areas interact and whether these interactions change depending on the specific behavioral context. LFP signals can provide insights into the neural processes underpinning various cognitive functions and actions^22,23,24^. We used long stretches of LFP recordings (7-10 minutes) and identified facial movement epochs using manual scoring, and an automatic algorithm based on optic flow measurements (see Methods). Functional connectivity was assessed using pairwise phase consistency (PPC) between the LFPs of the recorded areas, quantifying the distribution of phase differences^25^. The advantage of using PPC versus other connectivity measurements is that it remains reliable even with few trials, and it is less prone to errors that occur when the consistency of phase differences correlates with the strength of the signal^25^. This method assumes that neural synchronization between two areas will show a distribution of phase differences centered around a mean value. The analysis first examines differences in functional connectivity patterns between periods of facial stillness (rest) and movement. We found clear differences in the functional connectivity between the moving and the still face. Overall, during emotional behaviors, beta-range (13-40 Hz) connectivity between lateral and medial motor areas (M1 and M3) increased during movement. The M3-PMv connectivity in the beta range, however, showed behavior-specific patterns: it increased during lipsmacks (LS) but decreased during threats (T). For voluntary movements (chews, C), the medial-to-lateral connectivity shifted toward lower frequencies, showing increased theta (5-7 Hz) and alpha (8-12 Hz) connectivity while beta connectivity diminished during movement execution. Regarding lateral motor areas connectivity, both emotional and voluntary movements showed increased M1-PMv beta band connectivity during movement compared to rest, with voluntary movements additionally exhibiting enhanced theta and alpha connectivity (for details see Table S1, cluster-based nonparametric statistics with dependent samples t-test, p <= 0.05).

While the above analysis revealed general patterns of functional connectivity that point to movement-related modulation of interactions between these areas, our main analysis focused on understanding the movement-specific functional interactions between the sensorimotor face areas, as well as characterizing the directional flow of information within the network during different types of facial movements, specifically targeting the movement execution period, which enhances our ability to accurately define distinct actions. We first calculated functional interactions between medial and lateral motor areas during threats (T - red) and lipsmacks (LS - yellow), representing emotionally based gestures with opposite valence (Fig. 3A-C). In general, synchrony appears to be frequency-dependent and consistently stronger during threats compared to lipsmacks. Statistical analyses revealed that threat-related interareal synchrony was significantly stronger than lipsmack-related synchrony, mostly in the low frequency range (thick shades in Fig. 3A-C; cluster-based nonparametric statistics with dependent samples t-test, p <= 0.05, see Table S2) but not in the high-frequency range (i.e., beta range). Interestingly, the functional interactions between the medial (M3) and lateral (M1 or PMv) motor systems during emotional facial movements (depicted in Fig. 3A-B) were comparable in properties but weaker in magnitude compared to the interactions within the lateral subnetwork (M1 and PMV in Fig. 3C). In both cases, the synchrony during threats was stronger than during lipsmacks in the low-frequency range (<20 Hz), but not in the beta range.

**Figure 3.**
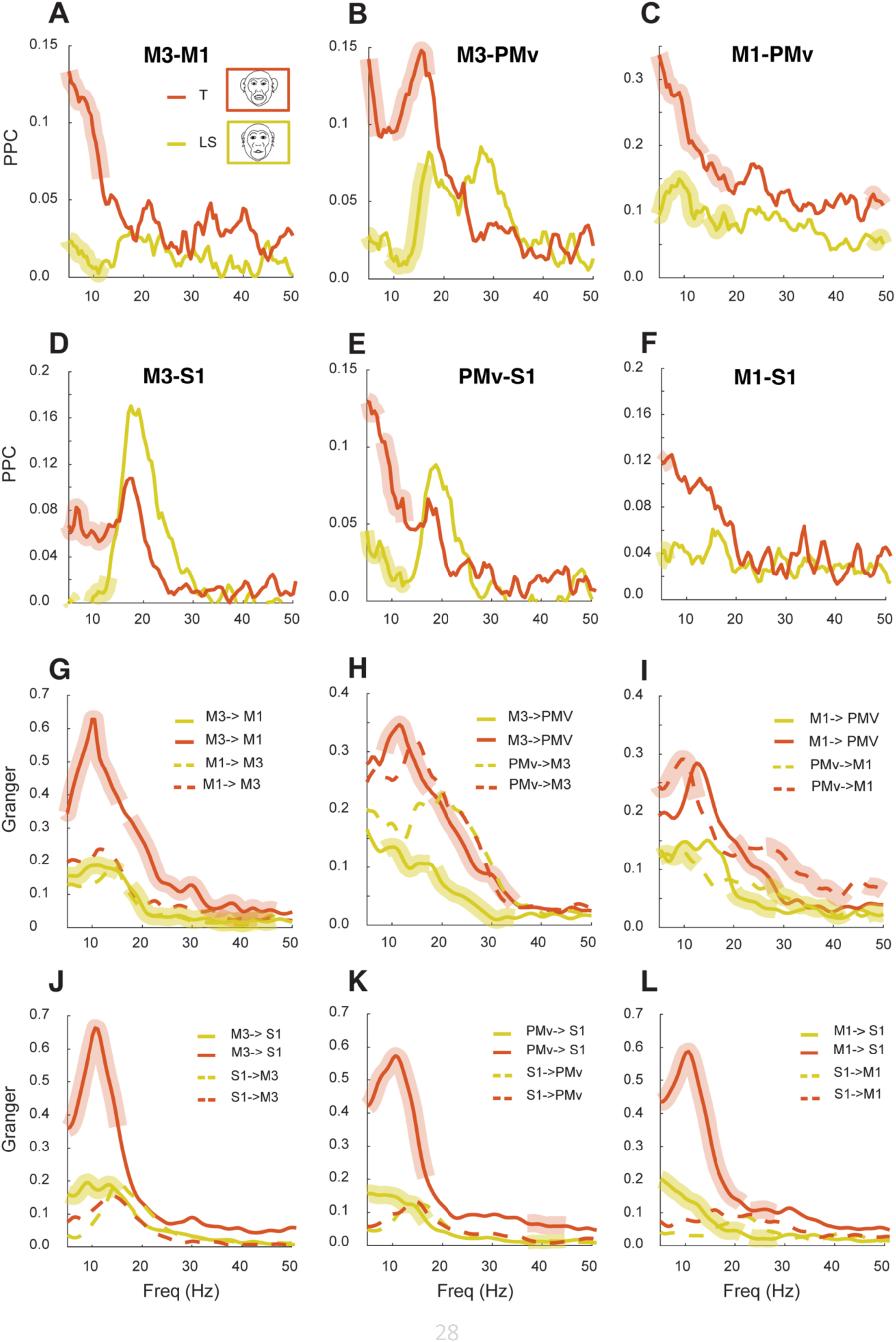
Functional interactions between facial motor and sensorimotor cortices during the production of emotional facial movements. **A-F** Pairwise phase consistency (PPC) between M3 and M1 (medial and lateral motor areas), M3 and PMv (medial and lateral motor areas), M1 and PMv (lateral motor areas), M3 and S1 (medial motor and primary somatosensory area), PMv and S1 (lateral and primary somatosensory area), M1 and S1 (lateral motor and primary somatosensory area) during the production of threats (T: red solid lines, n=1200) and lipsmacks (LS: yellow solid lines, n=1200). In Panels A-F thicker, semi-transparent lines atop the solid lines denote regions of statistical differences between both facial expressions (p≤0.05). **G-L** Granger causality for pairs of areas: M3-M1, M3-PMv, M1-PMv, M3-S1, PMv-S1, M1-S1 respectively. Solid lines depict a particular directionality of Granger causality (e.g., M3 to M1), while dashed lines depict the opposite direction (e.g. M1 to M3). Thicker semi-transparent lines indicate Granger causality values statistically different between the facial movements.

Beyond motor system interactions, we investigated how primary somatosensory cortex (S1) functionally connects with motor areas during different emotional gestures, providing insight into sensorimotor integration. We observed beta synchrony between S1 and both medial (M3) and lateral motor areas, with a clear broad peak in the beta frequency range for functional connectivity between S1 and PMv (Fig 3D-E) during threats and lipsmacks. Similar to patterns within the motor network, coupling between S1 and motor areas was consistently stronger during threats than during lipsmacks (Fig 3D-F). This gesture-specific synchrony pattern suggests functional differences beyond the distinct kinematics characterizing these two gestures.

Next, we compared the neural activity during emotional facial movements with that observed during voluntary facial movements (specifically, chewing movements). We found increased synchronization both between the lateral motor cortices (M1 and PMv), and between the medial and lateral motor cortices (M3 and M1, M3 and PMv) during chewing compared to rest (Table S1). Further analysis revealed consistent coupling within the lateral system (M1 and PMv) during both voluntary and emotional movements. While no significant differences were detected at high frequencies (>20 Hz) both threats and chews showed significant higher synchronization compared to lipsmacks at low frequencies (Fig. S2C, S2F; Table S2). Moreover, during voluntary chewing, the M3 area showed synchronization with M1(S2A, S2D) and PMv (S2B, S2E) mainly at low frequencies (4-8 Hz). In contrast, beta range synchronization was not prominent. In general, emotional movements elicited stronger alpha and beta coupling across the medial and lateral systems, while voluntary chewing movements relied more on low-frequency coupling, illustrating frequency-specific functional interactions during movement execution (Fig. S2A-F).

In summary, the lateral motor system (M1 and PMv) maintains consistent beta-frequency coupling during all movement types, with evident movement-specific differences occurring in the low frequency range (theta and alpha range), where both threats and chews showed stronger coupling than lipsmacks. The medial-to-lateral connectivity (M3 to M1 & PMv) shows broader frequency-range variations that depend on the ongoing movement type: emotional movements engage primarily alpha and beta range, while voluntary chews show reduced synchronization in these bands and predominantly operate at lower frequencies (<10 Hz, Table1).

### Directional Information within the Face Sensorimotor Network

Our findings have revealed a dynamic sensorimotor network where frequency coupling varies with the type of facial movement. However, the synchrony measurement we used (PPC) cannot identify directionality of interactions. For this purpose, we employed Granger Causality analysis, which measures the extent to which activity in one area (e.g., variable X) can significantly predict subsequent activity in another area (e.g., variable Y) thereby hinting at causal relations (or “information flow”) between the two regions^26,27^. We used Granger causality to reveal differences in inter-areal information flow during different facial movements, with a particular emphasis on information flow between medial and lateral areas of the facial motor system (M3 and M1 & PMv). Using this approach, we found bidirectional information flow between medial and lateral motor areas, with the magnitude of the flow varying according to the facial expression. When comparing threats to lipsmacks, information flow from medial to lateral motor areas was stronger during threats, while flow in the opposite direction remained similar between both behaviors (Fig. 3G-H). Interestingly, the interactions with S1 were substantially more asymmetric, showing a consistently stronger flow of information from motor regions to S1 than in the reverse direction (Fig. 3J-L).

During voluntary chewing movements, we observed slightly lower information flow from M3 to the lateral motor areas (M1 and PMv) when compared with threat-related emotional movements (Fig. S2J-K). Comparison between chewing and lipsmack movements revealed target-specific patterns. While M3-to-PMv information flow was similar between both movement types (Fig. S2H), M3 exerted a significantly stronger influence on M1 during chewing than during lipsmacks (Fig. S2G). Within the lateral system, higher information flow from M1 to PMv was detected during threats (9-15.5 Hz) compared to voluntary chewing. In contrast, information flow from PMv to M1 was similar between both movement types (Fig. S2L).

As shown in Figure 4, the network analysis reveals that motor areas show largely symmetric information flow (except for M3-to-M1 during threats Fig. 4A, or chews S2J), while motor-to-somatosensory information flow is predominantly unidirectional (Fig. 4A-B). While inter-areal synchrony was stronger within the lateral motor areas and weaker for medial-lateral motor interactions, information flow remains reliable across both medial and lateral regions, indicating continuous information exchange through the facial motor system (Fig. 4B). The pattern of differential synchrony strength coupled with reliable information exchange demonstrates how this sensorimotor network maintains flexible functional interactions. These interactions can be dynamically modulated to coordinate neural activity patterns underlying different facial gestures.

**Figure 4.**
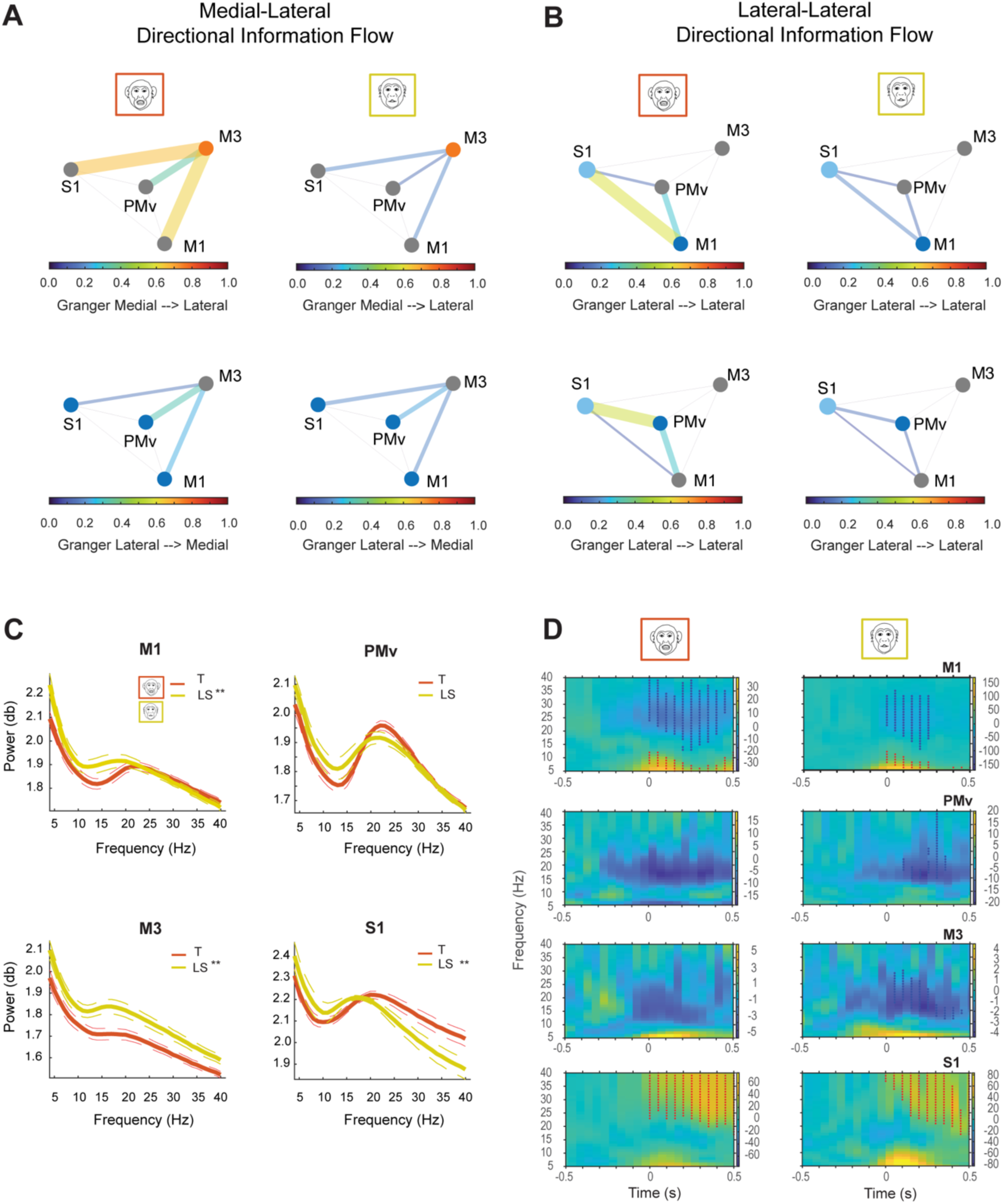
Summary of Network analysis during facial expressions, and local oscillatory activity within motor face and S1 areas. **A-B.** Summary network diagram depicting the Granger’s Causality maximum peak between each pair of motor areas and S1 during threats (left panel) and lipsmacks (right panel). Color and thickness of the line indicate the flow’s maximum magnitude for each direction. (**A**) Medial-Lateral Directional Information Flow: The upper panel depicts the flow from medial (M3, orange node) to lateral motor and S1 areas, while the lower panel shows flow from lateral areas (M1-PMV-S1, blue nodes) to medial motor area (M3). (**B**) Lateral-Lateral Directional Information Flow: Upper panel depicts flow from M1 to lateral motor areas (PMv and S1), while the lower panel shows flow from area PMv (blue node) to M1 and S1. (**C**) Raw power around the movement onset (±1s) for threats (T: red) and lip-smacks (LS: yellow) in M1, PMv, M3 and S1. Alpha (α = 8-12Hz) and beta (β = 13-40 Hz) activity was modulated across all behaviors. Asterisks indicate significant power differences in α and β ranges between threats and LS (p≤0.01, Kruskal-Wallis test corrected for multiple comparisons, n=2 monkeys, T=432, LS=292). (**D**) Time frequency representations (TFR, 5-40 Hz) normalized to pre-movement epoch for threats and lipsmacks in M1, PMv, M3, and S1. Time zero indicates movement onset. Asterisks show significant clusters (p≤0.05, cluster-corrected nonparametric randomization test) comparing pre-movement (-0.75 to - 0.25s) versus movement epochs (0.0 to 0.5s). Red asterisks denote activity increases, blue asterisks indicate decreases relative to baseline.

### Facial Movement-Specific Local Power Dynamics

After identifying distinct connectivity and directionality patterns during emotional and voluntary facial movements, we investigated how local neural activity in each network area relates to facial gestures, focusing on oscillatory power modulations that reflect the local neural population activity^22^. To investigate this, we analyzed the region-specific LFP dynamics.

Figure 4C and Fig S4A-C, illustrate modulation in the raw local power across the different frequency bands (alpha: 8-12 Hz, beta:13-40 Hz and gamma 40-120 Hz) in lateral (M1 and PMv), medial (M3) and somatosensory (S1) areas during different facial expressions, analyzed from 1s pre- to 1s post-movement onset. Significant differences in the power between behaviors were confirmed by the Kruskal-Wallis test (p<=0.01, Table S4-S5), with each behavior showing distinct patterns of power modulation across regions.

To assess the temporal pattern of the LFP signals in different areas and varying facial movements we normalized the power of the ongoing oscillations during the movement period against the preparatory period (see Methods). In motor areas we found a ubiquitous pattern of beta-suppression as has been identified during voluntary movements of the upper limb^5,28^. Nonetheless the details of the suppression dynamics were movement and area-dependent (Fig. 4D). For instance, in M1 the beta suppression was locked to movement onset and lasted longer during threats than lipsmacks. In PMv, beta suppression showed higher amplitude, ramped up a bit before the movement onset, and reached a maximum power during movement. In M3, suppression was prominent, more pronounced and longer for lipsmacks than for threats. Voluntary chew movements also evoked beta suppressions with dynamics slightly different from emotional movements (for details see Fig. S3).

In addition, we found in both M1 and M3 an increase in gamma power (>40 Hz), during emotional and voluntary movements (Fig. S4E-G). In S1, beta suppression was modest, reaching significance only during chewing movements (Fig. S3), while movement-related increases in alpha, high beta (> 20 Hz), and gamma power, were dominant across all movement types.

In summary, our findings reveal area-specific oscillatory patterns reflecting most likely distinct neural computations within each region. Activity in both medial and lateral facial motor areas showed only subtle differentiation between emotional and voluntary movements. In contrast, we observed robust interareal synchronization based on consistent phase relationships predominantly in alpha/beta bands that varied systematically across different facial gestures. These synchronized interactions between medial area M3, lateral areas (M1, PMv), and S1 occurred with frequency-specific coupling patterns unique to each facial gesture.

The magnitude and directionality of information flow within this network dynamically reconfigures depending on the type of facial movement.

## Discussion

In this study, we identified functional interactions between medial and lateral facial motor areas and the primary somatosensory cortex during facial expressions and voluntary movements in non-human primates, with facial expressions representing behaviors that are fundamental for social communication. Our results demonstrate a distributed cortical sensorimotor network with functional overlap between the previously proposed separate emotional (medial) and voluntary (lateral) circuits at the cortical level. This finding, at the cortical level, challenges the classical two-pathway model based on stroke patient studies^12,14^ and brain imaging research^29,30^ which proposes separate control circuits for emotional and voluntary facial movements. We instead found modulations in the local activity of both medial and lateral facial motor areas for both types of movements. While there was only subtle differentiation in local activity (measured as site-specific LFP power) between movement types, we identified specific frequency-based coupling among these regions. This suggests that facial expressions could emerge from a broad sensorimotor network with dynamic and flexible connectivity patterns that adapt based on the movement type.

The prominence of these functional interactions was particularly evident during emotional facial expressions, where medial and lateral motor cortices showed synchronized activity despite their distinct local oscillatory signatures. This integrated sensorimotor network perspective aligns with recent human studies^31^ showing that lateral motor areas receive emotional movement-related information, as evidenced by M1 activation during both voluntary speech production and emotional movements like smiling. Crucially, our ICMS evoked interareal connectivity results provide evidence for this network architecture, by revealing connectivity between medial and lateral facial motor areas. The consistency of interareal interactions across both methodologies – neural synchronization and electrically evoked responses – validates our results about functional interactions within this motor network.

Neural communication among populations is mediated, at least in part, by their synchronous activity^23,32^, which is fundamental for the dynamic coordination of distributed neural activity in local and extended networks underlying sensorimotor processes ^33,34,35,36^. Information processing between distant neural assemblies depends on the strength of coherent activity through synchronous oscillations^36^, with long range interactions involving a broader spectrum of frequency bands including theta, alpha and beta frequencies^37^.

Our results align with and extend these fundamental principles of neural communication, revealing specific patterns of synchrony between the neural populations of medial (M3) and lateral motor areas (M1 and PMv), as well as with the somatosensory cortex (S1) in the alpha and beta range, during facial expressions, extending into the theta range for voluntary chewing. The synchrony between areas, measured as phase relationships between the oscillations, was independent of changes in local power, and both the degree of synchrony and inter-area influence varied with the type of facial movement.

Alpha synchrony typically reflects the inhibition of task-irrelevant information^38,39^, crucial for executive functions ^36^. The patterns we observed in the face sensorimotor network align with this gating role. We found differential alpha coupling between lipsmacks and threats in key sensorimotor areas (medial-lateral: M3-PMV, M3-M1, M3-S1; lateral: M1-PMV, PMV- S1; Table S2) suggesting expression-specific synchrony modulations. This supports alpha’s role in gating information based on context and attention^39^. The modulations in alpha synchrony across facial expressions likely regulate information flow during facial movements by suppressing irrelevant inputs while maintaining attention on motor behavior.

Complementing the role of alpha synchrony, beta synchrony plays a crucial role in the large-scale coupling of sensorimotor information^40^. Beta band coherency is ideally suited for flexibly and dynamically forming neural assemblies^41^, and long-distance inter-area communication^37^. Beta oscillations can also be context-specific, reflecting current task rules or decisions^42,43^. In our case, shifts in the beta frequency peaks across different facial expressions (Fig 3A-B medial-lateral connectivity M3-M1, M3-PMv; Table 1) may contain information about the decision to perform specific facial movements in particular social contexts^43,44^. The significance of beta synchrony extends beyond our specific findings. Previous research has demonstrated beta synchrony’s crucial role in synchronizing large-scale cortical networks in sensorimotor^45^ and social decision-making tasks^46^. The occurrence of beta oscillations in the face motor system, with expression-specific frequency shifts, reinforces beta’s role as a rhythm for sensorimotor network synchronization across different effector systems, while suggesting these variations may reflect the selection process underlying specific facial movements.

**Table 1.**
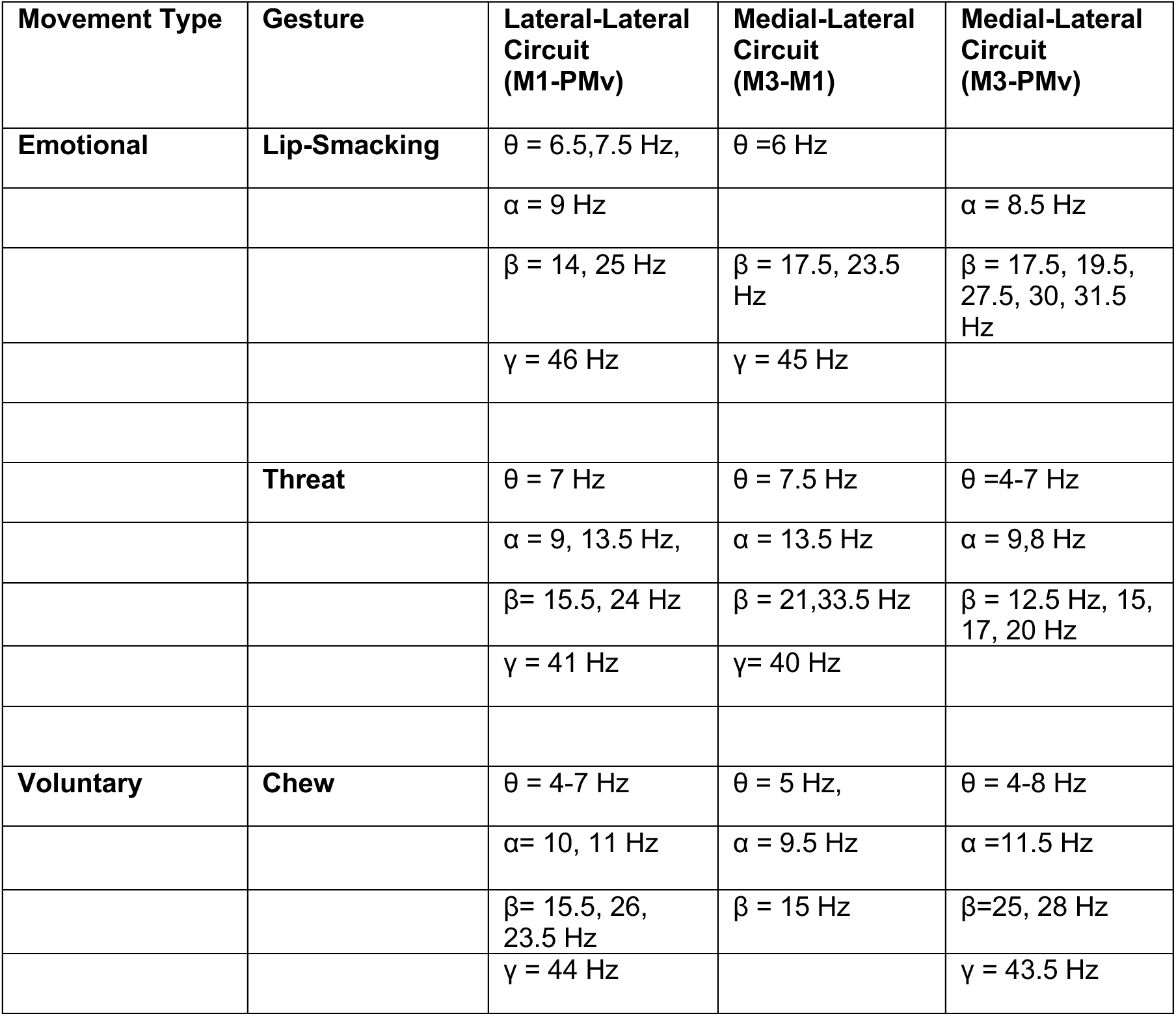
Peak Coupling Frequency Among Motor Circuits

| **Movement Type** | **Gesture** | **Lateral-Lateral Circuit (M1-PMv)** | **Medial-Lateral Circuit (M3-M1)** | **Medial-Lateral Circuit (M3-PMv)** |
| --- | --- | --- | --- | --- |
| **Emotional** | **Lip-Smacking** | $\theta = 6.5, 7.5 \text{ Hz}$, | $\theta = 6 \text{ Hz}$ | |
| | | $\alpha = 9 \text{ Hz}$ | | $\alpha = 8.5 \text{ Hz}$ |
| | | $\beta = 14, 25 \text{ Hz}$ | $\beta = 17.5, 23.5 \text{ Hz}$ | $\beta = 17.5, 19.5, 27.5, 30, 31.5 \text{ Hz}$ |
| | | $\gamma = 46 \text{ Hz}$ | $\gamma = 45 \text{ Hz}$ | |
| | **Threat** | $\theta = 7 \text{ Hz}$ | $\theta = 7.5 \text{ Hz}$ | $\theta = 4-7 \text{ Hz}$ |
| | | $\alpha = 9, 13.5 \text{ Hz}$, | $\alpha = 13.5 \text{ Hz}$ | $\alpha = 9, 8 \text{ Hz}$ |
| | | $\beta = 15.5, 24 \text{ Hz}$ | $\beta = 21, 33.5 \text{ Hz}$ | $\beta = 12.5 \text{ Hz}, 15, 17, 20 \text{ Hz}$ |
| | | $\gamma = 41 \text{ Hz}$ | $\gamma = 40 \text{ Hz}$ | |
| **Voluntary** | **Chew** | $\theta = 4-7 \text{ Hz}$ | $\theta = 5 \text{ Hz}$, | $\theta = 4-8 \text{ Hz}$ |
| | | $\alpha = 10, 11 \text{ Hz}$ | $\alpha = 9.5 \text{ Hz}$ | $\alpha = 11.5 \text{ Hz}$ |
| | | $\beta = 15.5, 26, 23.5 \text{ Hz}$ | $\beta = 15 \text{ Hz}$ | $\beta = 25, 28 \text{ Hz}$ |
| | | $\gamma = 44 \text{ Hz}$ | | $\gamma = 43.5 \text{ Hz}$ |

In summary, our data confirm that phase coupling is highly specific to gesture type, frequency, and the areas engaged in the coupling. The distinction between emotional and voluntary facial movements is frequency dependent: coupling at low frequencies (theta, alpha) shows larger differences within the emotional movements category (threat> lipsmack), while coupling in the beta range between medial and lateral areas (M3 - PMV) and between these areas and S1 reveals a clear categorical distinction where both emotional expressions exhibit stronger coupling than voluntary chewing. This suggests low frequencies encode information specific to each gesture, whereas beta coupling may specifically encode aspects related to the production of emotional movements.

To understand how this synchrony-mediated coordination occurs, we examined information flow using Granger Causality analysis, which quantifies the extent to which activity in one area significantly predicts subsequent activity in another area. Results showed continuous bidirectional exchange between medial and lateral motor regions, as well as among lateral regions, indicating dynamic information flow during facial expressions. We observed behavior-specific asymmetries, such as enhanced information flow from medial area M3 to lateral motor area M1 during threats. This asymmetry might reflect the unique demands of threat displays, which require sustained, non-affiliative movements that may depend more heavily on M3 control, unlike the affiliative and rhythmic patterns of lipsmacks in which the M3- to-M1 information flow was attenuated. Importantly, while we categorized lipsmacks as emotional movements, other studies suggest that they may be evolutionary precursors of human speech, potentially incorporating both emotional and voluntary control components^47^.

This dual nature, along with the opposing valence between threats and lipsmacks, may contribute to the observed asymmetric information flow pattern from medial to lateral areas. In contrast, PMv, showed symmetric information flow patterns between medial and lateral areas during both emotional movements, suggesting it could function as a central hub facilitating bidirectional information exchange (Fig. 3H-I)^48^. The primary somatosensory cortex exhibited asymmetric communication patterns, receiving more information from motor areas than it sent back. While this incoming flow might represent information about planned or ongoing movement^49^, further studies are needed to test this hypothesis.

Our results demonstrate that information flow within the face sensorimotor network exhibits varying patterns of functional symmetries and asymmetries, resulting in different degrees of influence between the interconnected areas. One could speculate about the existence of a flexible functional hierarchy within this network shaped by the kind of facial movement, similar to what has been proposed for the lateral forelimb motor cortices in mice^50^. Such a flexible, context-dependent, switch in hierarchy may explain the discrepancy between the extensive medial-to-lateral interactions found in this study and the specific emotional vs. voluntary impairments reported in patients suffering from medial vs. lateral lesions^12^.

In summary, our findings reveal a dynamic cortical sensorimotor network with flexible connectivity that adapts based on the behavioral context and facial movement type. Intracortical stimulation showed that medial and lateral areas exert significant functional impact on each other, while simultaneous LFP recordings confirmed these regions interact primarily in alpha and beta frequency ranges during facial expressions. Rather than separate motor cortical circuits for emotional and voluntary movements, our data suggest that facial control relies on coordinated inter-areal interactions within an extensive sensorimotor network. This finding challenges the hypothesis of independent cortical medial/lateral streams for facial motor control. This network likely enhances signal coordination to the facial nucleus by integrating information across brain regions. Importantly, cortical integration does not exclude the possibility of downstream segregation.

Beyond the sensorimotor network itself, a key question for future research is how this network for facial expression production interfaces with a broader social network comprising the STS and limbic structures^1,51^. The STS, recognized as the third visual pathway^52^, provides dynamic social cues such as facial expressions, eye gaze, and body movements - while limbic structures (insula, orbitofrontal cortex, and amygdala) encode emotional and reward information. Understanding how these systems interface through the cingulate and premotor areas ^20, 53–56^ is crucial for explaining how value, emotion, and social context influence facial motor output decisions. Based on known anatomical connections, we propose that social and emotional information flows from these regions—with the STS detecting dynamic social signals and limbic structures computing emotional valence and reward value – converging in cingulate and premotor areas where these inputs are integrated into the motor network to guide the decision about which facial expression to produce.

Understanding this integrated network architecture has important clinical implications. Facial palsy affects over 60% of stroke patients, with deficits often persisting beyond 40 days^57^, causing significant functional and aesthetic impairments^58^. While traditional predictive models focus on lesion characteristics, these have not translated into effective therapeutic interventions. Our findings offer a different contribution: we identify the cortical facial motor network (M1-PMV, M3-PMV, M3-M1 extending to S1) as a potential therapeutic target for neuromodulation-based rehabilitation strategies. Specifically, PMv showed symmetric information flow patterns between both medial and lateral areas, identifying it as a potential hub facilitating bidirectional information exchange (Fig. 3H-I, Fig. S2H, K).

These findings suggest specific application for post-stroke facial palsy rehabilitation using network-targeted neuromodulation (TMS, tDCS), which has shown promise in motor recovery by inducing spike-timing-dependent plasticity ^59–63^. While bihemispheric approaches may yield different results for facial versus limb motor control given the automaticity of symmetric facial muscle activation, our results particularly highlight PMv as a target for intra-hemispheric neuromodulation therapy. Targeting PMv with neuromodulation, with the aim of enhancing both medial (PMv-M3) and lateral (PMv-M1) cortical connectivity upstream to the frequently injured M1, would be both a technically feasible and well-motivated hypothesis.

Finally, we acknowledge the putative overlap between motor and premotor cortical representations and descending outputs to laryngeal motor neurons controlling vocalizations ^64^, highlighting the need for future experiments to elucidate the neural mechanisms underlying co-occurring facial expressions and vocal output during social communication. Understanding both the dynamics of this cortical network during facial expressions alone versus when accompanied by vocalizations, and the temporal dynamics between these behaviors, is fundamental to understanding how human speech could have evolved.

## Materials and Methods

All animal housing, care, and experimental procedures complied with the National Institutes of Health Guide for Care and Use of Laboratory Animals and were approved by the Institutional Animal Care and Use Committees of the Rockefeller University (protocol numbers 21104-H USDA) and Weill Cornell Medical College (protocol number 2010-0029).

### Subjects

We studied two male monkeys (Macaca fascicularis (M1) and Macaca mulatta (M2)) that were group -or pair housed before and during the experiments.

A cranial implant composed of MR-compatible acrylic cements, anchored with MR-compatible ceramic screws was implanted in each monkey following standard surgical procedures, as well as standard anesthetic, aseptic and postoperative treatment protocols. A custom-designed MR-compatible headpost was fixed to each implant.

### fMRI Behavioral Task

Monkeys were trained to sit in a sphinx position and maintain passive fixation on a dot at the center of a screen for 2-4 seconds to receive fluid reward while blocks of pictures or videos were presented, interleaved with baseline periods during which only the fixation dot appeared.

During imagining, the subjects sat in an MRI compatible NHP chair with the head fixed at isocenter. Gaze position was tracked at 120 Hz, with an MRI compatible eye-tracker. All behavioral and stimulus display parameters were controlled with Presentation Software (Neurobehavioral Systems). During the functional scans, video stimuli were projected at 60 Hz resolution on a back-projection screen placed 35 cm in front of the subject’s eyes. Monkeys performed a social-interaction free-viewing task as described in Shepherd & Freiwald^20^, and an oro-facial motor localizer in which sparse, intermittent fluid reward was delivered during a passive fixation task^65^.

Facial movements were tracked at 15 Hz using an MRI-compatible infrared video camera (MRC), whose acquisition was synchronized with the first TR of each scan run. The onset and offset of the facial emotional (threats, lipsmacks), voluntary (chew, drinking) and other (yawn) movements was determined offline by two independent observers, manually annotated and cross-validated.

### fMRI Acquisition

Monkey2 was previously fMRI-mapped as described in Shepherd & Freiwald^20^. Monkey1 was scanned and fMRI mapped in a similar fashion. All MRI data were acquired in a 3T Siemens Prisma scanner, while monkeys were in sphinx position. Functional imaging was acquired using custom-designed surface 8 channel receiver radiofrequency coils and a horizontal single loop transmit coil (L. Wald, MGH, Martinos Center for Biomedical Imagining) in echoplanar imagining sequences (EPI: TR 2.25 s, TE=17 ms, Flip angle 79 degrees, 1.2 mm^3^ isotropic voxels, FOV=96 mm). Matrix size= 80 X 80 X 45, horizontal interleaved slices, 2X GRAPA acceleration. Volumes per run ranged from 168 volumes per run.

Right before each scanning session, Molday ION (monocrystalline iron oxide nanoparticles) was injected into the saphenous vein below the knee to increase the contrast-to-noise ratio (CNR^66^). The dose ranged from 9 mg/kg on an initial scan day to 6 mg/kg on subsequent scanning days.

Anatomical images were collected from each subject in a separate session while subjects were anesthetized (ketamine 5-8 mg/kg, isoflurane 0.5-2.0%, and dexmedetomidine 0.008- 0.02mg/kg), and placed on an MR-compatible stereotactic frame. Images were acquired using a customized 1-channel receive coil (L. Wald, MGH, Martinos Center for Biomedical Imagining). Anatomical images were based on the averaged of 6 repetitions of a T1-weighted magnetization-prepared rapid gradient echo (MPRAGE) sequence (FOV 128 mm, voxel size 0.5 X 0.5 X 0.5 mm). A computed tomography (CT scan) of the head & neck was acquired on the same day, to provide high anatomical resolution of the skull and skin.

### fMRI Analysis and Face Sensorimotor Cortex Localization

Imagining analysis was done with FreeSurfer and FS-FAST (v6.0), using customized MATLAB and Linux-shell scripts as described in Shepherd & Freiwald ^20^. Raw image volumes were 2D (slice-wise) motion and time corrected (AFNI, 3dAllineate, version AFNI_2011_12_21_1014), aligned to high-resolution anatomical T1, unwarped (JIP Analysis Toolbox, v3.1), smoothed (2 mm Gaussian FWHM) and masked. The first four volumes of each functional run were excluded from further analysis.

We constructed two GLM models to localize the sensory and motor face cortical areas. The onset times of the manually scored facial expressions were convolved with the MION HRF to generate a primary regressor of interest that will result in a “face expression activity map”. In the second GLM model the times at which juice-was delivered, were used to create an “orofacial motor activity map”.

Average signal intensity maps for each contrast were computed with FS-FAST function selxavg3-sess. The MION HRF was modeled as a wide gamma function with a steep initial slope (delta = 0; tau = 8000; alpha = 0.3; dt = 1 ms) and discretized to the length of the TR (2.25 s), in order to be used as a regressor in the GLM. In both models, nuisance regressors included 2^nd^ order polynomial for drift, and the top 3 head motion PCs obtained from 2D slice-wise motion correction (3DAllieniate). The resulting voxel-wise beta-weight maps were not further corrected for multiple comparisons.

Significant voxels in both maps were identified, with a conjunction analysis using logical ANDs, taking the least significant *p-value* from each contrast entered in the conjunction. The conjunction facial expression-orofacial motor map was used for surgical targeting. This reliably identified voxels primarily correlated with oro-facial movement, and not body movements during the social-interaction free viewing task.

Brain coordinates were transformed from RAS space to stereotaxic coordinates using the Saleem-Logothetis atlas^67^, to verify target locations were within the intended cortical areas for surgical implantation. Transformation aligned AC to +21mm from the interaural line, with coordinates calculated as distances from reference points: AP from AC (Y-axis), ML from midline (X-axis), and DV from ear bar zero (Z-axis).

Monkey 1 (M. fascicularis):

● Reference: AC [-0.18, 24.18, 19.28], PC [-0.18, 15.13, 18.94], EBZ [-0.52, 7.12, 12.01] DV adjusted with 1.14 scaling factor and + 4.71mm offset for species differences
● Targets [RAS → AP/ML/DV mm]: S1:[21.49, 19.36, 26.46] → [+16.18/+21.67/+21.18] M1 medial: [20.79, 22.71, 26.98] → [+19.53/+20.97/+21.78] M1 lateral: [22.62, 22.69, 24.78] → [+19.51/+22.80/+ 19.27]PMv: [19.01, 30.32, 26.90] →[+27.14/+19/+21.68] M3: [3.99, 34.05, 30.98] → [+30.87/+4.17/+26.34]

Monkey 2 (M. mulatta):

● Reference: AC [-3.69, -3.01, 24.67], PC [-3.69, -16.91, 23.43], EBZ [-3.65, -24.38, 10.65]
● Targets [RAS → AP/ML/DV mm]: S1: [17.28, -8.88, 37.86] → [+15.13/+20.97/+27.21] M1: [17.25, -5.08, 37.86] → [+18.93/+20.94/+27.21] PMv: [18.43, 3.37, 33.38] → [+27.38/+22.12/+22.73] M3: [0.18, 7.78, 39.47] → [+31.79/+3.87/+28.82]

### fMRI-Guided Electrophysiological Recording sites

We used CORTEXPLORER SCI, a commercially available integrated functional/structural MRI-based stereotaxic planning and intra-surgical tracking approach (cortEXplore GmbH, Linz Austria) to target the position of 6-8 FMAs (floating microarrays, Microprobes: https://microprobes.com/products/multichannel-arrays/array-comparison-chart) based on the fMRI facial expression-orofacial motor map, described above. We targeted the face representation in the following areas based on the fMRI map: primary somatosensory cortex (S1), primary motor cortex (M1), ventro-lateral premotor cortex (PMv) and rostral cingulate motor area (M3). All arrays were implanted in the right hemisphere of both animals (Fig. 1C and Fig. S1).

For each subject, we aligned all functional maps and a high-resolution CT to the anatomical MRI, creating a project with the images of the skin, skull, brain and functional maps. Thus, the surgical planning occurred in the subjects’ native space. For each target, we defined a corresponding entry point on the cortical surface; together these two points define a 3D trajectory for the array implantation. Each FMA included electrodes of varying length to maximize gray matter neural activity recordings. Intraoperative, we registered the surgical subject with its radiological data in CORTEXPLORER SCI, under general anesthesia and sterile conditions. This process creates a spatial relationship between the 3D virtual image space and the 3D physical subject.

We registered the subject in two steps – first with seven fiducial markers on the subject’s acrylic implant, and second, with a few hundred points collected from the real subject’s surface, in which the probe scans the surface of the surgical subject, registering it to its corresponding virtual surface. For each subject, the registration error was <200 microns RMS. Based on the target coordinates in each animal, we performed craniotomies and durotomies to facilitate the implantation of the FMAs. Each FMA was loaded onto the optically tracked probe, aligned with a predefined trajectory, and advanced slowly into the cortex for implantation. In all cases the 3D angular error between the actual and predefined trajectories was <1 degree.

### Neural Recording Behavioral Task

We used a randomized block design where each recording day was divided into 7–10 minute blocks, with behavioral and electrophysiological data recorded from head-fixed monkeys in different behavioral conditions. Block order was randomized to control for time and attentional effects, with a minimum of 3 different block types recorded daily: rest (baseline/control condition), voluntary chewing/ingestive movements, and emotional facial expressions for social communication including threats (aggressive gestures) and lipsmacks (affiliative gestures). During rest blocks, animals remained calm with minimal movement. During chew blocks, animals were intermittently fed fruit pieces. Facial expressions were elicited using a battery of stimuli including conspecific face stimuli from social interaction task^20^, conspecific movie stimuli, monkey avatar stimuli, and live interactions with conspecifics and humans. Animals were previously trained on fixation tasks to ensure attention to stimuli. During recordings, the reward tube was removed to prevent mouth occlusion and disambiguate social expressions from ingestive movements. While facial expression blocks were generally dominated by one expression type, some datasets contained both threats and lipsmacks within the same block.

All visual stimuli were presented, and behavior controlled using MonkeyLogic (v2.2), running on a Windows computer system which sent the triggers to the TDT (Tucker-Davis Technologies) data-acquisition system via an analog and digital input/output card PCIe-6343. Subjects viewed visual stimuli on a 56 x 24 cm (1024 x 768 resolution) screen at ∼58 cm from the eyes, with a refresh rate of 60 Hz. The conspecific movie stimuli were recorded with a GoPro7 camera attached to the monkeys’ home cages. We extracted ten-second clips of monkeys engaging in social behaviors e.g. grooming and playing, and non-social behaviors e.g. eating, drinking, and idling. Videos contained one or two monkeys, either male or female, and a subset of these monkeys were familiar to the subject being recorded. Each clip was phase-scrambled (to control for low-level motion, color and luminance intensity). One stimulus set shown during a single recording run consisted of 7-10 individual clips and their matched scrambles presented in a pseudo-random order.

For a subset of recordings, subjects viewed an interactive monkey avatar face. The avatar’s facial movements were controlled by real-time tracking of the experimenter’s face using an Optitrack system connected to a PC running Unity. The experimenter was in a separate room from the subject.

Every trial began with a variable period of required visual fixation, after which a video stimulus appeared; the subject was able to freely move his eyes to explore the videos as long as his gaze stayed within a virtual window the size of the frame of the video and additional 2 degrees surrounding it. No reward was given during these runs, and the reward tube was removed.

### Face and Eye Movements Recordings

Facial movements and expressions were recorded using a FLIR BFS-US-13Y3 (http://softwareservices.flir.com/BFS-U3-13Y3/latest/Model/spec.html, monkey 1) camera at a video sampling rate of 70 Hz and a Flex 13 Optitrack system at 120Hz (https://optitrack.com/cameras/flex-13/, monkey 2) mounted ∼70 cm away from the subject’s face. All videos were black & white. Videos were captured by a dedicated Windows computer running Spinview (FLIR) or Motive (Optitrack). An additional Motive camera was used to monitor lower body movements during the experiments. Crucially, we did not observe during the experiment a systematic co-occurrence between any of the three facial movements studied and body movements. Body movements occurred randomly and were temporally uncorrelated with the onset of the facial movements. Video acquisition was synchronized with a TDT system using a TTL pulse. Eye movements were recorded using an ISCAN system and synchronized with the TDT (1.017 kHz) and Monkey Logic system.

#### Electrophysiological Recordings

Neural activity was recorded simultaneously using FMA arrays with 36 channels per array (FMA, Microprobes Inc) with spacing of 400 µm, and 4 x 1.8 mm dimensions. We recorded a total of 256 channels (M1) and 192 (M2) of 6-8 individual arrays implanted in the face representation of the following areas: S1, M1, PMv and the rostral motor cingulate (M3). In each array the electrode length was customized for each area to maximize gray matter activity recordings. In monkey 1, we used for analysis the arrays localized in the medial regions for M1 and PMv since the signal to noise ratio, quality and modulation of the signal was better than for their lateral counterparts. Electrode lengths ranged from 1.2-2.8 mm in all sensory-motor areas (S1, M1, PMv), except for M3 (5-9 mm) which is a deeper structure. In addition, the lengths of the electrodes were staggered and increased towards the sulcus. Recording electrode impedances were ∼ 0.5 MOhm (n=28), microstimulation electrode impedance ∼ 10K and less than 10 K for ground (n=2) and reference (n=2) electrodes.

#### Data Acquisition

Neural activity was recorded at full bandwidth with a sampling frequency of 24 kHz, and a resolution of 16 bits using a Tucker-Davis Technologies (TDT) system RZ2 Biosignal Processor. Neural data and all behaviors were synchronously stored to disk together with the face videos and eye movements. Raw recordings were filtered offline to obtain LFP (1kHz).

#### Functional Connectivity Paradigm

In monkey 1, the paradigm consisted in applying a (250 µA, a train of single biphasic pulses, each pulse duration=0.4 ms at a frequency of 3 Hz (333 ms inter-pulse interval) delivered for ∼2min) through each of the stimulation electrodes. Current acquisition was done at 24 kHz. Neural activity was stored as described above. Each array contained four low impedance (∼10K) electrodes evenly spaced across the array, and 28 recording electrodes. Crucially, stimulation was delivered while the animal sat quietly without making facial or body movements. Using this current protocol, we did not evoke any facial or body movements. These parameters were specifically chosen to evoke neural responses in the receiver area rather than overt movements.

We first pre-processed and cleaned the data by selecting per array the channels without noise, using this channel selection we averaged their LFPs, and then analyzed the averaged evoked LFP across repetitions (n=50 repetitions, 32 or less channels per array, n=3 days). Per recording day, normalization was achieved by calculating a z-score per recording electrode and array. The z-score was determined by the voltage of the given electrode minus the average voltage across all selected electrodes of the array divided by their standard deviation. All z-scores per array for all given days were averaged to get a grand-average day where the connectivity measurements were measured.

Connectivity was measured by quantifying the amplitude of the evoked Local Field Potential (eLFP) response in the receiver areas following current delivery from a sender area. We measured the absolute difference between successive peak and trough amplitudes (Δ|Peak-Trough) in the eLFP. To avoid measuring stimulation artifacts, we searched for peaks and troughs starting 6 ms after stimulation onset in all receiving areas, when artifactual deflections had ended. These connectivity measurements were then represented in a confusion matrix.

#### Behavioral Analysis

Facial expressions, voluntary chewing movements and other facial movements, were manually scored by 3 human observers for one monkey. The onset of the facial movements was determined by the time at which the subject’s mouth started to move preceded by one second of mouth stillness. For the other monkey, we used for half of the sessions a customized python based automatic detector for mouth movements, based on the optic flow of the recorded video. This automatic scoring method was based on the average magnitude of optical flow around the animal’s mouth in the recorded video. For each session, a Gaussian mixture model estimating the average optical flow magnitude around the animal’s mouth, was used to cluster the frames in two categories. One category involved small or no mouth movements at all, while the other category involved large movements. The period of a facial movement was defined as a large movement period (>30 ms), preceded and followed by non-movement or small movement period longer than 1 second. For validation, the automatic onsets were cross validated with manual scoring onsets.

To further validate our movement detection and provide an objective framework for categorizing movement types, we used a semi-supervised approach to parse behavior as well as neural activity using PCA. Specifically, we extracted segments containing Deep Lab Cut marker trajectories^68^ and LFP signals from 500 ms before to 1000 ms after each manually scored movement onset, which served as input to the PCA. In summary, we could separate the DLC markers and the neural activity into three categories corresponding to the following facial movements: threat, lipsmacks and chews, which validates our labelling. The first three principal components accounted for more than 80% of behavioral variance and were primarily associated with mouth and nostril markers. Combined with our fMRI localization and targeting of the recorded areas (see methods above) this suggests that a considerable portion of the neural variance within our recorded regions relates to mouth and surrounding facial movements.

#### Spectral Analysis

In both monkeys, the LFP data were bandpass filtered from 1-256 Hz and preprocessed to form a data structure containing all voltage recordings per channel within each array, the sampling rate, and all trials with their corresponding behavioral events. All voltages were analyzed in µV.

We used custom-built MATLAB R2023a code (the MathWorks Inc., Natick, Massachusetts, USA) and Fieldtrip toolbox (v20170101)^69^ to analyze the data. We analyzed a total of 13(T), 13(C) & 7 (LS) recording sessions. A band-stop filter in monkey 2 was applied to remove the line noise at 60 Hz and harmonics. Artifacts, dead channels, and trials containing artifacts related to movement or electronic interference were removed based on visual inspection of the data.

The voltage was re-referenced per array using the low impedance electrodes distributed across the array. Using a fast Fourier transform approach, we computed time frequency representations (TFR) of the power (5-120Hz). For the range of 5-40 Hz, we used an adaptive sliding time window 4 cycles long (Dt=4/f) multiplied with a Hanning taper. For higher frequencies (40-120Hz), we used multitapers sleepians in a 0.2 s window with +/- 10 Hz of smoothing. The windows moved in .05 s steps, from 1s before the movement onset to 1s after the movement execution. Per type of facial movement condition i.e., threat, lipsmack or chew we averaged over trials within each recording session and then normalized by an absolute baseline subtraction using a 0.75 s window before movement onset. For visualization purposes, a grand average was computed over the recording sites (per area). In the case of the raw power, a similar analysis was performed, except for the normalization. Statistical analysis – see below, were computed using the raw power.

#### Connectivity analysis

We measured functional connectivity between the pairs of recorded areas by computing the pairwise phase consistency (PPC) measurement as implemented in Fieldtrip ^25,69^. The PPC is an unbiased and consistent estimate of how well two signals generated by two sources show a consistent phase relationship in a particular frequency band (5-50 Hz). The advantage of using PPC versus other connectivity measurements is: 1) it remains reliable even with few trials, and 2) it is less prone to errors that occur when the consistency of phase differences correlates with the strength of the signal^25^.

In the case of the connectivity analysis, we focused on the movement execution period, to maximize trial numbers and ensure certainty about the nature of the ongoing facial movement type. We divided each full movement period into 0.5 second epochs, with each epoch serving as an individual trial for that behavior. Movements lasting less than 0.5 seconds (which constituted a minor proportion of all movements) were excluded from the analysis. To avoid sample biases when comparing functional connectivity and Granger causality across behaviors, we equalized trial numbers for each comparison. For each comparison, trials were first equalized within individual session pairs. The final sample sizes are depicted in table 2. Importantly, we ensure equal trial numbers between sessions for each behavior comparison – for example, 50 trials from a lip-smacking session were analyzed alongside 50 trials from the corresponding threat session. This approach allowed us to make robust statistical comparisons across different facial movement types.

**Table 2.** Number of trials for power and time frequency analysis.

| Facial Movement Type | Number of Trials |
| --- | --- |
| Threats | 432 |
| Lipsmacks | 292 |
| Chew | 415 |

Data from 2 monkeys. See figures 4, S3, S4.

**Table 3.**
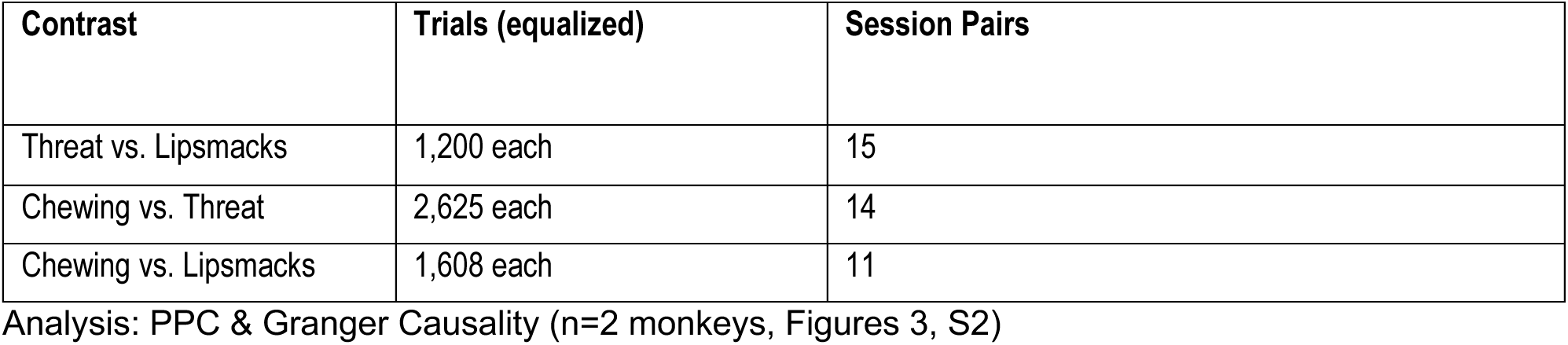
Trial counts for connectivity contrasts.

As a first approach, we calculated the connectivity (PPC) of the network when the face was at rest, i.e. not moving or planning to move and compared it with the connectivity when the face was moving. Statistical differences within this contrast were assessed by a cluster-based nonparametric randomization^69^, with a dependent samples t-test as statistic (see methods Statistical Analysis)

We used bivariate, nonparametric Granger Causality as implemented in Fieldtrip during the movement period to investigate the directionality of the connectivity^70^ during the movement epoch.

### Statistical Analysis

Differences in the average raw power across behaviors were calculated using a Kruskal-Wallis test, corrected for multiple comparisons (p<=0.01). The asterisks in Fig. 4A-D and Fig. S4A-D denotes significant differences in the mean power between threats and other behaviors. However, all the performed comparisons are depicted in Table S4.

In the case of the Time Frequency Representation (TFR), statistical differences in the power were assessed comparing a window of 0.5 s before the movement onset (-0.75s to -0.25s) versus 0.5 s time window during the movement execution (0s to 0.5s).

We applied a cluster-based nonparametric randomization^69^, with a dependent samples t-test to establish whether there were significant differences between two conditions (e.g. pre-movement vs movement epoch, or lipsmacks vs threats) in terms of power, PPC or Granger causality. By clustering neighboring samples (time-frequency points that show the same effect), this statistical test deals with the multiple-comparisons problem while accounting for the dependency of the data. All samples for which this t-value exceeded a prior threshold (uncorrected p<=0.05) were selected and subsequently clustered based on temporal-spectral adjacency. The sum of the t-values within a cluster was used as the cluster-level statistic. This was computed within all recording sessions (separately per area). By randomizing the data across the two conditions and recalculating the test statistic 1000 times, we obtained a null distribution of maximum cluster t-values to evaluate the statistic of the data (p<=0.05).

### Editorial Assistance

We used the AI-based language model, ChatGPT v4 by OpenAI and Claude 4.5 Sonnet, for English language editing to enhance manuscript clarity and readability. This comprises suggestions for sentence restructuring and phrasing improvements. Neither of them contributed to the scientific content or interpretation of our results. The final version of this manuscript was thoroughly reviewed and revised by all the authors, who are responsible for all content.

## Acknowledgments

Our sincere thanks to Gonzalez A., Yin L., Halibart B., Aboharb F., Landi S., Serene S. and the Freiwald Lab. We are very thankful to Shepherd S. V. for imagining analysis of monkey 2, and to Schaffelhofer S. for the development of neurosurgical tracking technologies (cortEXplore, Linz Austria) and the fMRI targeting of monkey 2, which was supported by Shepherd’s contributions. Thanks to the RU Veterinary Staff for surgical support. We thank Hudspeth A.J. for being a source of scientific inspiration, and the Hudspeth lab for their support, as well as Haegens S., Astafurov A. and Vergara J. for thoughtful discussions on data analysis and encouragement.

This work was supported by National Institute Of Neurological Disorders And Stroke of the National Institutes of Health under Award Number R01NS110901, the Charles H. Revson and Leon Levy Foundation Postdoctoral Fellowships (V. Y.); MSTP grant from NIGMS NIH # T32GM007739 to the Weill Cornell/Rockefeller/Sloan Kettering Tri-Institutional MD-PhD Program and NIMH/NIH Ruth L. Kirschstein NRSA #F30MH122157 (I.G.R); Austrian Science Fund (FWF) Erwin Schrödinger Fellowship J4580 (R.E); The Price Family Center for the Social Brain (WAF); US-Israel Binational Science Foundation (BSF) #2023321 (YP, WAF). The content is solely the responsibility of the authors and does not necessarily represent the official views of the NIH and the ONR.

## Supplementary Information

**Fig. S1.**
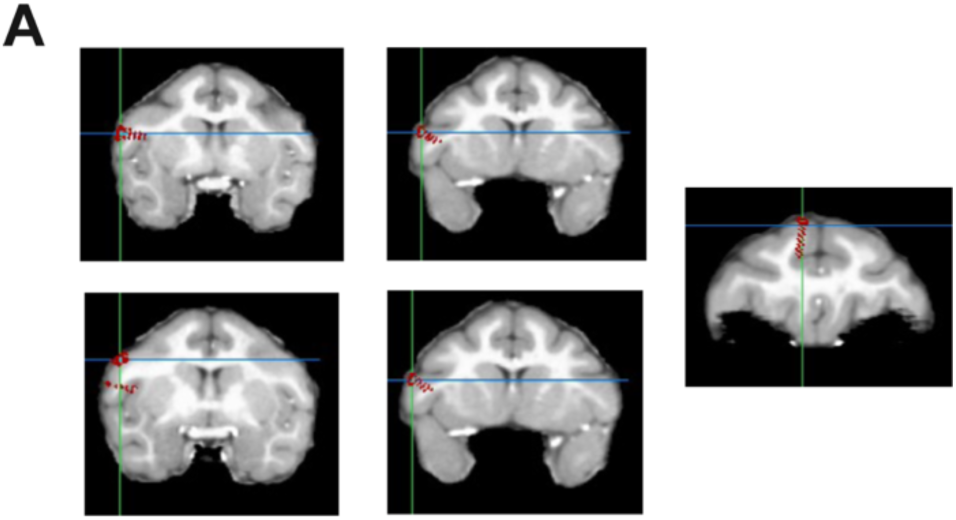
Anatomical locations of the arrays in the monkey’s brain. **(A)** Structural MRI (T1, coronal plane) and overlay showing electrode targeting for each array in the second monkey. Floating microarrays were placed over the primary somatosensory (S1) and motor (M1) cortex, ventrolateral premotor (PMv) and cingulate motor cortex (M3).

**Fig. S2.**
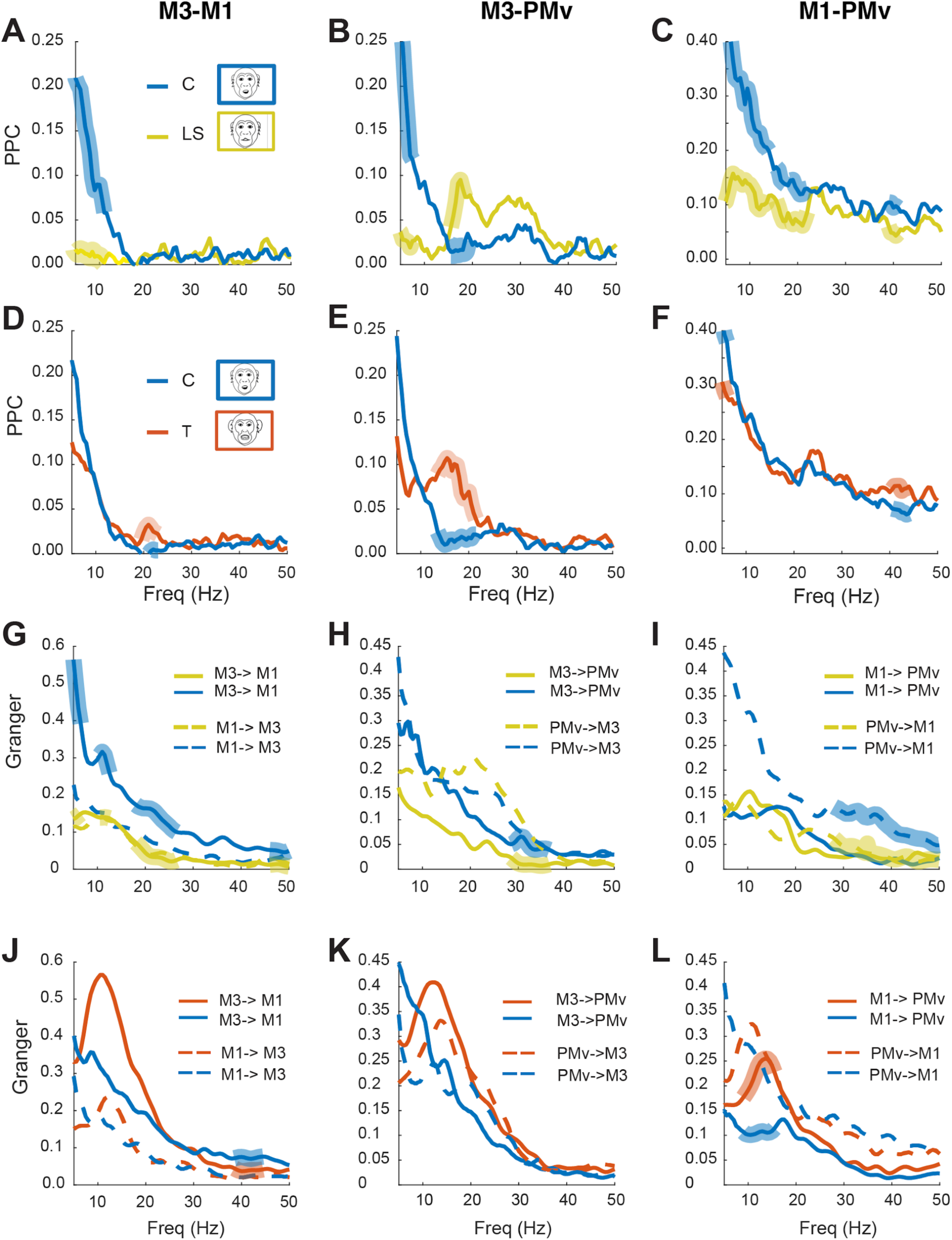
Functional interactions between medial and lateral motor areas during emotional and voluntary movements. **A-C.** Pairwise phase consistency (PPC) between M3-M1, M3-PMv, and M1-PMv during the production of lipsmacks (LS: yellow) versus voluntary chew movements (C: blue). **D-F.** Same PPC measures during threats (T: red) versus voluntary chews (blue). **G-L.** Granger causality for pairs of areas: M3-M1, M3-PMv, and M1-PMv, from left to right. Solid lines depict a particular directional flow (e.g. M3 to M1), while dashed lines depict the opposite direction, (e.g. M1 to M3). Thicker semi-transparent lines indicate statistically different values between movements (p ≤ 0.05). For details see the main text.

**Fig. S3.**
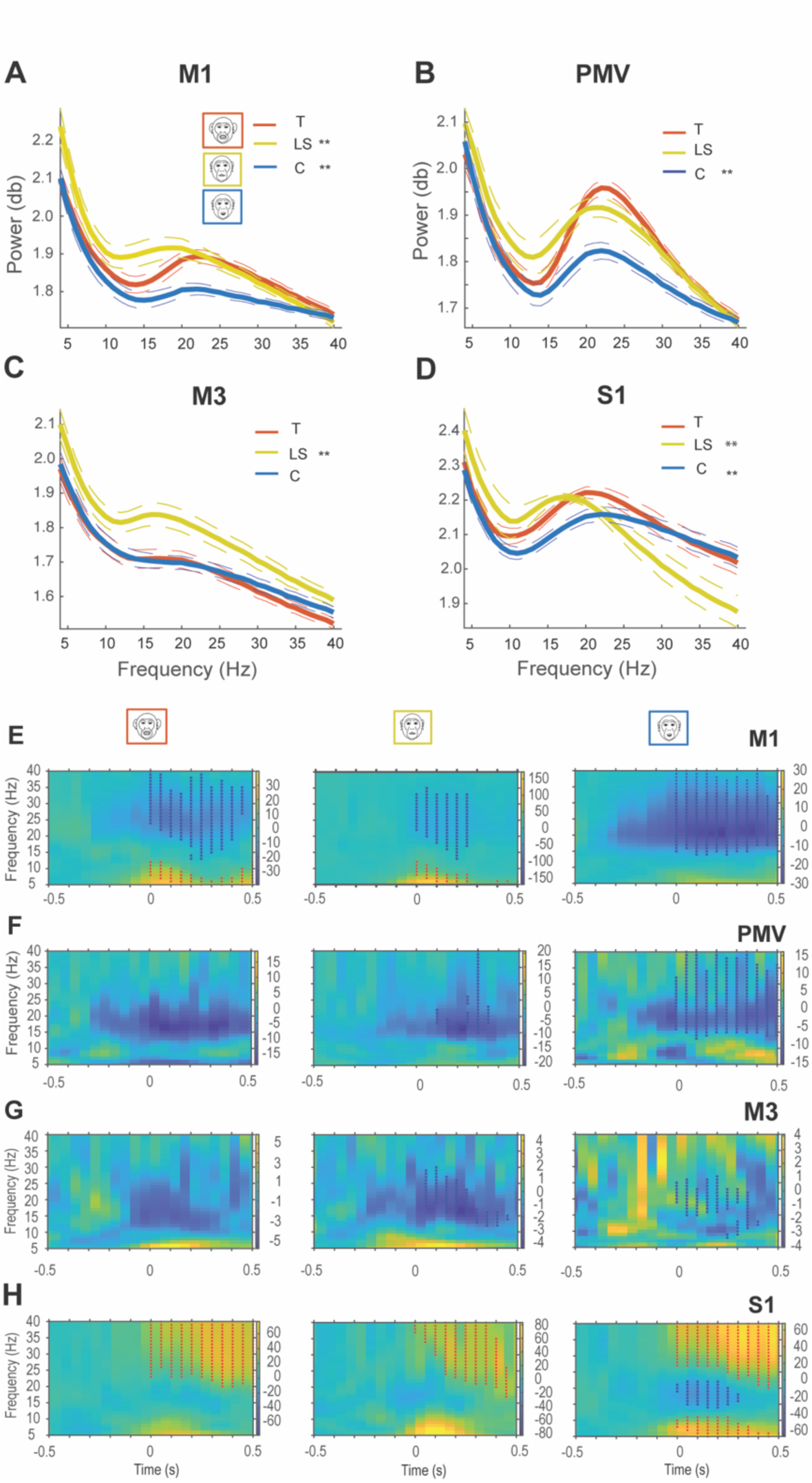
Oscillatory activity within the sensory and motor face areas for emotional and voluntary facial movements. **A-D** Raw power around the movement onset (±1s) for threats (T: red), lip-smacks (LS: yellow) and chews (C: blue) in M1 (**A**), PMv (**B**), M3 (**C**) and S1(D). Alpha (α = 8-12Hz) and beta (β = 13-40 Hz) activity was modulated across all behaviors. Asterisks indicate significant power differences in α and β ranges between threats and other behaviors (p≤0.01, Kruskal-Wallis test corrected for multiple comparisons, df=2, n=2 monkeys, T=432, LS=292, C=415). **E-H** Time frequency representations (TFR, 5-40 Hz) normalized to pre-movement epoch for threats, lipsmacks and chews in M1 (**E**), PMv (**F**), M3 (**G**), and S1 (**H**). Time zero indicates movement onset. Asterisks show significant clusters (p≤0.05, cluster-corrected nonparametric randomization test) comparing pre-movement (-0.75 to -0.25s) versus movement epochs (0.0 to 0.5s). Red asterisks denote activity increases, blue asterisks indicate decreases relative to baseline.

**Fig. S4.**
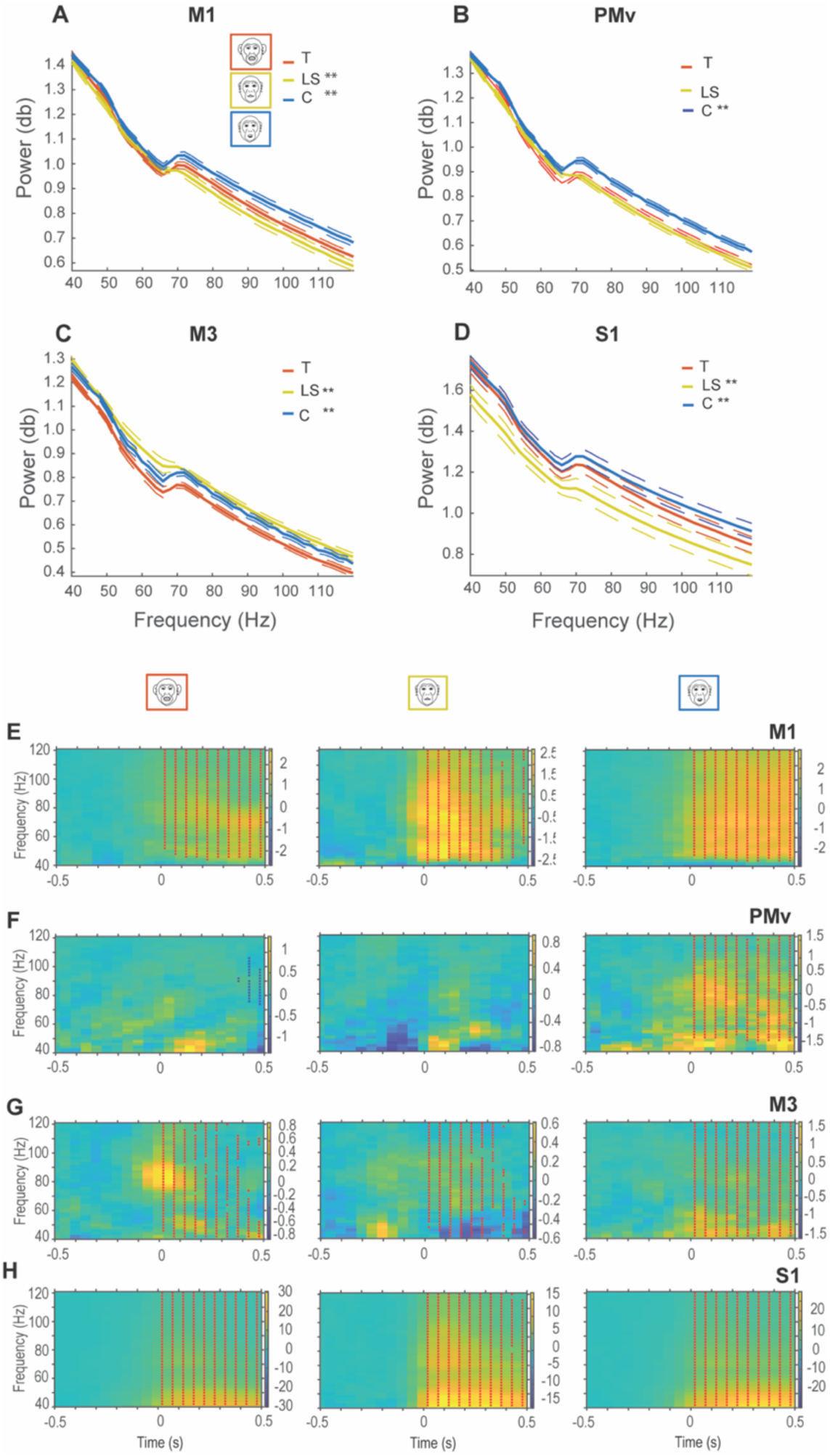
Oscillatory activity (40-120Hz) within sensory and motor face areas during emotional and voluntary facial movements. **A-D** Raw power around movement onset (±1s) for threats (T: red), lip-smacks (LS: yellow) and chews (C: blue) in: M1 (**A**), PMv (**B**), M3 (**C**) and S1 (**D**). Asterisks indicate significant power differences in gamma range (40-120 Hz) between threats and other behaviors (p≤0.01, Kruskal-Wallis test corrected for multiple comparisons, df=2, n=2 monkeys, T=432, LS=292, C=415). **E-H.** Time frequency representations normalized to pre-movement epoch (-0.75 to -0.25s) for threats, lipsmacks and chews in M1(**E**), PMv(**F**), M3(**G**) and S1(**H**) areas. Asterisks indicate significant clusters (p≤0.05, cluster-corrected nonparametric randomization test) comparing pre-movement (-0.75 to -0.25s) versus movement epoch (0.0 to 0.5s). Red asterisks denote power increases, blue asterisks decreases relative to baseline.

**Table S1.** PPC during Facial Movement *vs* Rest.

| Facial Movement | Area Pair | Frequency Range | pvalue | Permutations |
| --- | --- | --- | --- | --- |
| LS | M1-PMV | 18.5 - 20 Hz | $p \leq 0.05$ | 1000 |
| LS | M1-M3 | 16-18 Hz | $p \leq 0.05$ | 1000 |
| LS | M3-PMV | 31-33.5 Hz | $p \leq 0.05$ | 1000 |
| LS | M1-S1 | 40-40.5 Hz | $p=0.13$ | 1000 |
| LS | M3-S1 | 12-16.5 Hz | $p \leq 0.05$ | 1000 |
| LS | PMV-S1 | 14.5-17 Hz | $p \leq 0.05$ | 1000 |
| T | M1-PMV | 22-24.5 Hz | $p \leq 0.05$ | 1000 |
| T | M1-M3 | 28.5-29 Hz | $p=0.15$ | 1000 |
| T | M3-PMV | 19-21.5, 35-36.5 Hz | $p=0.07$ | 1000 |
| T | M1-S1 | 37.5-39 Hz | $p \leq 0.05$ | 1000 |
| T | M3-S1 | 12-15 Hz | $p \leq 0.05$ | 1000 |
| T | PMV-S1 | Tendency in $\beta$ range | $p>0.05$ | 1000 |
| C | M1-PMV | 4-19Hz<br>32-35 Hz | $p \leq 0.05$ | 1000 |
| C | M1-M3 | 4-11 Hz | $p \leq 0.05$ | 1000 |
| C | M3-PMV | 4-15.5 Hz<br>18-26.5 Hz | $p \leq 0.05$ | 1000 |
| C | M1-S1 | 30-32 Hz | $p \leq 0.05$ | 1000 |
| C | M3-S1 | 4-8 Hz<br>19.5-24 Hz | $p \leq 0.05$ | 1000 |
| C | PMV-S1 | 4-8 Hz<br>9-15 Hz<br>22.5-24.5 Hz | $p \leq 0.05$ | 1000 |

\*Facial Movement: LS (Lips smack); T(Threat); C(Chew) for this and tables.

**Table S2.** PPC Among Facial Movements.

| Facial Movements | Area Pair | Frequency Range | pvalue | Permutations |
| --- | --- | --- | --- | --- |
| T vs LS | M1-PMV | 4-13.5 Hz, 15-19 Hz, 47.5-50 Hz | $p \leq 0.03$, $p \leq 0.017$<br>$p \leq 0.035$ | 1000 |
| T vs LS | M3-M1 | 4-11 Hz | $p \leq 0.003$ | 1000 |
| T vs LS | M3-PMv | 4-7 Hz, 9.5-16.5 Hz | $p \leq 0.035$, $p \leq 0.01$ | 1000 |
| T vs LS | M1-S1 | 4-6.5 Hz | $p \leq 0.0350$ | 1000 |
| T vs LS | M3-S1 | 4-13 Hz | $p \leq 0.0009$ | 1000 |
| T vs LS | PMV-S1 | 4-7 Hz, 8-12.5 Hz | $p \leq 0.011$, $p \leq 0.020$ | 1000 |
| T vs C | M1-PMv | ----- | $p > 0.05$ | 1000 |
| T vs C | M3-M1 | 21.5-24.5Hz | $p \leq 0.0009$ | 1000 |
| T vs C | M3-PMv | 10.5-15 Hz, 20-26 Hz | $p \leq 0.007$, $p \leq 0.003$ | 1000 |
| LS vs C | M1-PMv | 4-22 Hz, 39-41 Hz | $p \leq 0.05$ | 1000 |
| LS vs C | M3-M1 | 4-11 Hz | $p \leq 0.05$ | 1000 |
| LS vs C | M3-PMv | 4-7.5 Hz, 15.5 - 18.5 Hz | $p \leq 0.05$ | 1000 |

**Table S3.** Granger Causality Among Facial Movements.

| Facial Movements | Direction | Frequency Range | pvalue | Permutations |
| --- | --- | --- | --- | --- |
| T vs LS | M1 to PMV | 20-28.5 Hz | $p \leq 0.011$ | 1000 |
| | PMV to M1 | 4-12.5 Hz, 26.5-42.5 Hz, 46.5-50 Hz | $p \leq 0.018$, $p \leq 0.007$, $p \leq 0.043$ | 1000 |
| T vs LS | M1 to M3 | 35-43Hz | $p \leq 0.008$ | 1000 |
| | M3 to M1 | 4-15.5 Hz, 17.5-46.5 Hz | $p \leq 0.009$, $p \leq 0.0009$ | 1000 |
| T vs LS | M3 to PMV | 8.5-17.5 Hz, 18.5-34.5 Hz | $p \leq 0.02$, $p \leq 0.007$ | 1000 |
| | PMV to M3 | ----- | $p > 0.05$ | 1000 |
| T vs LS | M1 to S1 | 4-21 Hz, 23-28 Hz | $p \leq 0.002$, $p \leq 0.0230$ | 1000 |
|  | S1 to M1 | ----- |  | 1000 |
| T vs LS | M3 to S1 | 4-15 Hz | $p \leq 0.0060$ | 1000 |
| | S1 to M3 | ----- | $p > 0.05$ | 1000 |
| T vs LS | PMV to S1 | 4-16.5 Hz, 37-44.5 Hz | $p \leq 0.0100$, $p \leq 0.0230$ | 1000 |
| | S1 to PMV | ----- | $p > 0.05$ | 1000 |
| T vs C | M1 to PMV | 9-15.5Hz | $p \leq 0.05$ | 1000 |
| | PMV to M1 | ----- | $p > 0.05$ | 1000 |
| T vs C | M1 to M3 | ----- | $p > 0.05$ | 1000 |
| | M3 to M1 | 38.5-44.5 Hz | $p \leq 0.05$ | 1000 |
| T vs C | M3 to PMV | ----- | $p > 0.05$ | 1000 |
| | PMV to M3 | ----- | $p > 0.05$ | 1000 |
| LS vs C | M1 to PMv | 4-22 Hz, 39-41 Hz | $p \leq 0.06$<br>$p \leq 0.05$ | 1000 |
| | PMv to M1 | 28-50 Hz | $p \leq 0.05$ | 1000 |
| LS vs C | M1 to M3 | ----- | $p > 0.05$ | 1000 |
| | M3 to M1 | 4-6.5 Hz, 10.5-12.5 Hz, 18-26.5 Hz, 46.5-50 Hz | $p \leq 0.05$ | 1000 |
| LS vs C | M3 to PMV | 28.5-36.5 Hz | $p \leq 0.05$ | 1000 |
| | PMv to M3 | ---- | $p \leq 0.05$ | 1000 |

**Table S4.** Power across Different Facial Movements (8-40Hz)

| Brain Area, Movements | Diff [95% CI] | pvalue |
| --- | --- | --- |
| S1, T-LS | 214.91 [117.63, 312.19] | p≤0.001 |
| S1, T-C | 177.03 [102.72, 251.35] | p≤0.001 |
| S1, LS-C | -37.88 [-134.62, 58.87] | p=0.629 |
| M1, T-LS | -182.86 [-280.14, -85.58] | p≤0.001 |
| M1, T-C | 312.85 [238.53, 387.16] | p≤0.001 |
| M1, LS-C | 495.71 [398.97, 592.45] | p≤0.001 |
| PMv, T-LS | -64.61 [-161.89, 32.67] | p=0.265 |
| PMv, T-C | 366.00 [291.68, 440.31] | p≤0.001 |
| PMv, LS-C | 430.61 [333.87, 527.35] | p≤0.001 |
| M3, T-LS | -578.57 [-675.85, -481.29] | p≤0.001 |
| M3, T-C | -54.03 [-128.34, 20.29] | p=0.204 |
| M3, LS-C | 524.54 [427.80, 621.29] | p≤0.001 |

**Table S5.** Power across Different Facial Movements (40-120Hz)

| **Brain Area, Movements** | **Diff [95% CI]** | **pvalue** |
| --- | --- | --- |
| S1, T-LS | 388.86 [295.39, 482.33] | $p \leq 0.001$ |
| S1, T-C | -116.27 [-187.67, -44.87] | $p \leq 0.001$ |
| S1, LS-C | -505.13 [-598.08, -412.18] | $p \leq 0.001$ |
| M1, T-LS | 179.80 [86.33, 273.26] | $p \leq 0.001$ |
| M1, T-C | -308.16 [-379.56, -236.75] | $p \leq 0.001$ |
| M1, LS-C | -487.95 [-580.90, -395.00] | $p \leq 0.001$ |
| PMv, T-LS | 69.72 [-23.74, 163.19] | $p = 0.187$ |
| PMv, T-C | -431.00 [-502.41, -359.60] | $p \leq 0.001$ |
| PMv, LS-C | -500.73 [-593.68, -407.78] | $p \leq 0.001$ |
| M3, T-LS | -601.64 [-695.10, -508.17] | $p \leq 0.001$ |
| M3, T-C | -388.80 [-460.20, -317.39] | $p \leq 0.001$ |
| M3, LS-C | 212.84 [119.89, 305.79] | $p \leq 0.001$ |

## References

1. R. Baez-Mendoza, Y. Vazquez, E. P. Mastrobattista, Z. M. Williams, Neuronal Circuits for Social Decision-Making and Their Clinical Implications. Front Neurosci 15, 720294 (2021).

2. M. Brecht, W. A. Freiwald, The many facets of facial interactions in mammals. Curr Opin Neurobiol 22, 259–266 (2012).

3. S. R. Hage, Language evolution in primates. Science 385, 713–714 (2024).

4. M. Davare, A. Kraskov, J. C. Rothwell, R. N. Lemon, Interactions between areas of the cortical grasping network. Curr Opin Neurobiol 21, 565–570 (2011).

5. B. E. Kilavik, M. Zaepffel, A. Brovelli, W. A. MacKay, A. Riehle, The ups and downs of beta oscillations in sensorimotor cortex. Exp Neurol 245, 15–26 (2013).

6. M. M. Churchland et al., Neural population dynamics during reaching. Nature 487, 51–56 (2012).

7. T. Ouchi et al., Mapping eye, arm, and reward information in frontal motor cortices using electrocorticography in non-human primates. bioRxiv 10.1101/2024.08.13.607846 (2024).

8. R. J. Morecraft, K. S. Stilwell-Morecraft, W. R. Rossing, The motor cortex and facial expression: new insights from neuroscience. Neurologist 10, 235–249 (2004).

9. R. M. Muri, Cortical control of facial expression. J Comp Neurol 524, 1578–1585 (2016).

10. U. Livneh, J. Resnik, Y. Shohat, R. Paz, Self-monitoring of social facial expressions in the primate amygdala and cingulate cortex. Proc Natl Acad Sci U S A 109, 18956–18961 (2012).

11. R. J. Morecraft, J. L. Louie, J. L. Herrick, K. S. Stilwell-Morecraft, Cortical innervation of the facial nucleus in the non-human primate: a new interpretation of the effects of stroke and related subtotal brain trauma on the muscles of facial expression. Brain 124, 176–208 (2001).

12. H. C. Hopf, W. Muller-Forell, N. J. Hopf, Localization of emotional and volitional facial paresis. Neurology 42, 1918–1923 (1992).

13. P. Cerrato et al., Emotional facial paresis in a patient with a lateral medullary infarction. Neurology 60, 723–724 (2003).

14. R. T. Ross, R. Mathiesen, Images in clinical medicine. Volitional and emotional supranuclear facial weakness. N Engl J Med 338, 1515 (1998).

15. K. F. Muakkassa, P. L. Strick, Frontal lobe inputs to primate motor cortex: evidence for four somatotopically organized ’premotor’ areas. Brain Res 177, 176–182 (1979).

16. R. J. Morecraft, G. W. Van Hoesen, Cingulate input to the primary and supplementary motor cortices in the rhesus monkey: evidence for somatotopy in areas 24c and 23c. J Comp Neurol 322, 471–489 (1992).

17. R. J. Morecraft, C. M. Schroeder, J. Keifer, Organization of face representation in the cingulate cortex of the rhesus monkey. Neuroreport 7, 1343–1348 (1996).

18. H. Tokuno, M. Takada, A. Nambu, M. Inase, Reevaluation of ipsilateral corticocortical inputs to the orofacial region of the primary motor cortex in the macaque monkey. J Comp Neurol 389, 34–48 (1997).

19. R. J. Morecraft, G. W. Van Hoesen, Frontal R. J. Morecraft, G. W. Van Hoesen, Frontal granular cortex input to the cingulate (M3), supplementary (M2) and primary (M1) motor cortices in the rhesus monkey. J Comp Neurol 337, 669–689 J Comp Neurol 337, 669-689 (1993).

20. S. V. Shepherd, W. A. Freiwald, Functional Networks for Social Communication in the Macaque Monkey. Neuron 99, 413–420 e413 (2018).

21. B. Ruszala, K. A. Mazurek, M. H. Schieber, Somatosensory cortex microstimulation modulates primary motor and ventral premotor cortex neurons with extensive spatial convergence and divergence. bioRxiv 10.1101/2023.08.05.552025 (2023).

22. S. Katzner et al., Local origin of field potentials in visual cortex. Neuron 61, 35–41 (2009).

23. B. Pesaran et al., Investigating large-scale brain dynamics using field potential recordings: analysis and interpretation. Nat Neurosci 21, 903–919 (2018).

24. C. Gallego-Carracedo, M. G. Perich, R. H. Chowdhury, L. E. Miller, J. A. Gallego, Local field potentials reflect cortical population dynamics in a region-specific and frequency-dependent manner. Elife 11 (2022).

25. M. Vinck, M. van Wingerden, T. Womelsdorf, P. Fries, C. M. Pennartz, The pairwise phase consistency: a bias-free measure of rhythmic neuronal synchronization. Neuroimage 51, 112–122 (2010).

26. L. Barnett, A. B. Barrett, A. K. Seth, Granger causality and transfer entropy are equivalent for Gaussian variables. Phys Rev Lett 103, 238701 (2009).

27. A. K. Seth, A. B. Barrett, L. Barnett, Granger causality analysis in neuroscience and neuroimaging. J Neurosci 35, 3293–3297 (2015).

28. J. Barone, H. E. Rossiter, Understanding the Role of Sensorimotor Beta Oscillations. Front Syst Neurosci 15, 655886 (2021).

29. M. Iwase et al., Neural substrates of human facial expression of pleasant emotion induced by comic films: a PET Study. Neuroimage 17, 758–768 (2002).

30. B. Wild et al., Humor and smiling: cortical regions selective for cognitive, affective, and volitional components. Neurology 66, 887–893 (2006).

31. M. Kern, S. Bert, O. Glanz, A. Schulze-Bonhage, T. Ball, Human motor cortex relies on sparse and action-specific activation during laughing, smiling and speech production. Commun Biol 2, 118 (2019).

32. A. M. Bastos, J. M. Schoffelen, A Tutorial Review of Functional Connectivity Analysis Methods and Their Interpretational Pitfalls. Front Syst Neurosci 9, 175 (2015).

33. P. J. Uhlhaas et al., Neural synchrony in cortical networks: history, concept and current status. Front Integr Neurosci 3, 17 (2009).

34. S. Haegens, J. Vergara, R. Rossi-Pool, L. Lemus, R. Romo, Beta oscillations reflect supramodal information during perceptual judgment. Proc Natl Acad Sci U S A 114, 13810–13815 (2017).

35. M. A. Hagan, B. Pesaran, Modulation of inhibitory communication coordinates looking and reaching. Nature 604, 708–713 (2022).

36. C. Beste, A. Munchau, C. Frings, Towards a systematization of brain oscillatory activity in actions. Commun Biol 6, 137 (2023).

37. N. Kopell, G. B. Ermentrout, M. A. Whittington, R. D. Traub, Gamma rhythms and beta rhythms have different synchronization properties. Proc Natl Acad Sci U S A 97, 1867–1872 (2000).

38. W. Klimesch, alpha-band oscillations, attention, and controlled access to stored information. Trends Cogn Sci 16, 606–617 (2012).

39. S. Haegens, V. Nacher, R. Luna, R. Romo, O. Jensen, alpha-Oscillations in the monkey sensorimotor network influence discrimination performance by rhythmical inhibition of neuronal spiking. Proc Natl Acad Sci U S A 108, 19377–19382 (2011).

40. M. Siegel, T. H. Donner, A. K. Engel, Spectral fingerprints of large-scale neuronal interactions. Nat Rev Neurosci 13, 121–134 (2012).

41. N. Kopell, M. A. Whittington, M. A. Kramer, Neuronal assembly dynamics in the beta1 frequency range permits short-term memory. Proc Natl Acad Sci U S A 108, 3779–3784 (2011).

42. B. Spitzer, S. Haegens, Beyond the Status Quo: A Role for Beta Oscillations in Endogenous Content (Re)Activation. eNeuro 4 (2017).

43. E. Rassi et al., Distinct beta frequencies reflect categorical decisions. Nat Commun 14, 2923 (2023).

44. C. Constantinidis, A. A. Ahmed, J. D. Wallis, A. P. Batista, Common Mechanisms of Learning in Motor and Cognitive Systems. J Neurosci 43, 7523–7529 (2023).

45. B. Pesaran, M. J. Nelson, R. A. Andersen, Free choice activates a decision circuit between frontal and parietal cortex. Nature 453, 406–409 (2008).

46. O. Dal Monte, C. C. J. Chu, N. A. Fagan, S. W. C. Chang, Specialized medial prefrontal-amygdala coordination in other-regarding decision preference. Nat Neurosci 23, 565–574 (2020).

47. A. A. Ghazanfar, D. Y. Takahashi, N. Mathur, W. T. Fitch, Cineradiography of monkey lip-smacking reveals putative precursors of speech dynamics. Curr Biol 22, 1176–1182 (2012).

48. M. Petrides, G. Cadoret, S. Mackey, Orofacial somatomotor responses in the macaque monkey homologue of Broca’s area. Nature 435, 1235–1238 (2005).

49. G. Ariani, J. A. Pruszynski, J. Diedrichsen, Motor planning brings human primary somatosensory cortex into action-specific preparatory states. Elife 11 (2022).

50. A. Saiki-Ishikawa, M. Agrios, S. Savya, A. Forrest A, H. Sroussi, S. Hsu, D. Basrai, F. Xu, A. Miri, Hierarchy in influence but not firing patterns among forelimb motor cortices (2025). bioRxiv doi: 10.1101/2023.09.23.559136.

51. W. Freiwald, B. Duchaine, G. Yovel, Face Processing Systems: From Neurons to Real-World Social Perception. Annu Rev Neurosci 39, 325–346 (2016).

52. D. Pitcher, L. G. Ungerleider, Evidence for a Third Visual Pathway Specialized for Social Perception. Trends Cogn Sci 25, 100–110 (2021).

53. P. F. Ferrari, M. Gerbella, G. Coude, S. Rozzi, Two different mirror neuron networks: The sensorimotor (hand) and limbic (face) pathways. Neuroscience 358, 300–315 (2017).

54. N. Dolensek, D. A. Gehrlach, A. S. Klein, N. Gogolla, Facial expressions of emotion states and their neuronal correlates in mice. Science 368, 89–94 (2020).

55. A. Jezzini et al., A shared neural network for emotional expression and perception: an anatomical study in the macaque monkey. Front Behav Neurosci 9, 243 (2015).

56. K. M. Gothard, The amygdalo-motor pathways and the control of facial expressions. Front Neurosci 8,43(2014).

57. G. F. Volk et al., Facial motor and non-motor disabilities in patients with central facial paresis: a prospective cohort study. J Neurol 266, 46–56 (2019).

58. P. Konecny, M. Elfmark, K. Urbanek, Facial paresis after stroke and its impact on patients’ facial movement and mental status. J Rehabil Med 43, 73–75 (2011).

59. S. Lefebvre et al., Neural substrates underlying stimulation-enhanced motor skill learning after stroke. Brain 138, 149–163 (2015).

60. F. Fiori, E. Chiappini, A. Avenanti, Enhanced action performance following TMS manipulation of associative plasticity in ventral premotor-motor pathway. Neuroimage 183, 847–858 (2018).

61. S. Lefebvre et al., Differences in high-definition transcranial direct current stimulation over the motor hotspot versus the premotor cortex on motor network excitability. Sci Rep 9, 17605 (2019).

62. Lefebvre, S., Jann, K., Schmiesing, A. et al. Differences in high-definition transcranial direct current stimulation over the motor hotspot versus the premotor cortex on motor network excitability. Sci Rep 9, 17605 (2019).

63. S. Turrini et al., Transcranial cortico-cortical paired associative stimulation (ccPAS) over ventral premotor-motor pathways enhances action performance and corticomotor excitability in young adults more than in elderly adults. Front Aging Neurosci 15, 1119508 (2023).

64. C. M. Cerkevich, J. A. Rathelot, P. L. Strick, Cortical basis for skilled vocalization. Proc Natl Acad Sci U S A 119, e2122345119 (2022).

65. Z. Yang, W. A. Freiwald, Encoding of dynamic facial information in the middle dorsal face area. Proc Natl Acad Sci U S A 120, e2212735120 (2023).

66. F. P. Leite et al., Repeated fMRI using iron oxide contrast agent in awake, behaving macaques at 3 Tesla. Neuroimage 16, 283–294 (2002).

67. Salem KS, Logothetis NK. A combined MRI and Histology Atlas of the Rhesus Monkey Brain in Stereotaxic Coordinates. Academic Press, London UK, 1st Ed, (2007).

68. 68. G. R. Ianni et al., Facial gestures are enacted via a cortical hierarchy of dynamic and stable codes. bioRxiv 10.1101/2025.03.03.641159 (2025).

69. R. Oostenveld, P. Fries, E. Maris, J. M. Schoffelen, FieldTrip: Open-source software for advanced analysis of MEG, EEG, and invasive electrophysiological data. Comput Intell Neurosci, 156869 (2011).

70. M. Dhamala, G. Rangarajan, M. Ding, Estimating Granger causality from fourier and wavelet transforms of time series data. Phys Rev Lett 100, 018701 (2008).

